# Individual variations in “Brain age” relate to early life factors more than to longitudinal brain change

**DOI:** 10.1101/2021.02.08.428915

**Authors:** D. Vidal-Piñeiro, Y. Wang, SK. Krogsrud, IK. Amlien, WFC. Baaré, D. Bartrés-Faz, L. Bertram, A.M. Brandmaier, CA. Drevon, S. Düzel, KP. Ebmeier, RN Henson, C. Junque, RA Kievit, S. Kühn, E. Leonardsen, U. Lindenberger, KS. Madsen, F. Magnussen, AM. Mowinckel, L. Nyberg, JM. Roe, B. Segura, SM. Smith, Ø. Sørensen, S. Suri, R. Westerhausen, A. Zalesky, E. Zsoldos, the Australian Imaging Biomarkers and Lifestyle flagship study of ageing, KB. Walhovd, AM. Fjell

## Abstract

*Brain age* is a widely used index for quantifying individuals’ brain health as deviation from a normative brain aging trajectory. Higher than expected *brain age* is thought partially to reflect above-average rate of brain aging. We explicitly tested this assumption in two large datasets and found no association between cross-sectional *brain age* and steeper brain decline measured longitudinally. Rather, *brain age* in adulthood was associated with early-life influences indexed by birth weight and polygenic scores. The results call for nuanced interpretations of cross-sectional indices of the aging brain and question their validity as markers of ongoing within-person changes of the aging brain. Longitudinal imaging data should be preferred whenever the goal is to understand individual change trajectories of brain and cognition in aging.

## Introduction

The concept of *brain age* is increasingly used to capture inter-individual differences in the integrity of the aging brain^1^. The biological age of the brain is estimated typically by applying machine learning to magnetic resonance imaging (MRI) data to predict chronological age. The difference between predicted *brain age* and actual chronological age (*brain age delta*) reflects the deviation from the expected norm and is often used to index brain health. *Brain age delta* has been related to brain, mental, and cognitive health and proved valuable in predicting outcomes such as mortality^1–3^. To different degrees, it is assumed that *brain age delta* reflects past and ongoing neurobiological aging processes^1,3–6^. Hence, it is common to interpret positive *brain age deltas* as reflecting a steeper rate of brain aging; often dubbed as accelerated aging^1,4,6^.

The assumption that *brain age delta* reflects an ongoing process of (faster or slower) neurobiological aging implies that there should be a relationship between cross-sectional and longitudinal estimates of *brain age*. Alternatively, deviation from the expected *brain age* could show lifelong stability and capture earlier genetic and environmental influences^3,7,8^. These perspectives offer fundamentally divergent interpretations of higher *brain age (delta)* in groups experiencing specific life events, brain disorders, and other medical problems. Here we tested whether *brain age* is related to accelerated brain aging, early-life factors, or a combination of both (**Fig. 1a**). If interindividual variations of *brain age* reflect variations in rates of ongoing brain aging, cross-sectional *brain age delta* should be positively associated with brain decline measured longitudinally. Here, we quantified individual brain change as the annual rate of change of *brain age delta (brain age delta*_*long*_). In addition, we also assessed brain change with a composite score of brain change and change in the different *raw* brain features. If early-life influences play a substantial role, one should observe a relationship between *brain age* and early factors - indexed here by birth weight and polygenic scores for *brain age* (PGS-BA) given evidence of lifelong effects of genetic effect on age-related phenotypes^9,10^ (**Fig. 1b**).

**Fig. 1.**
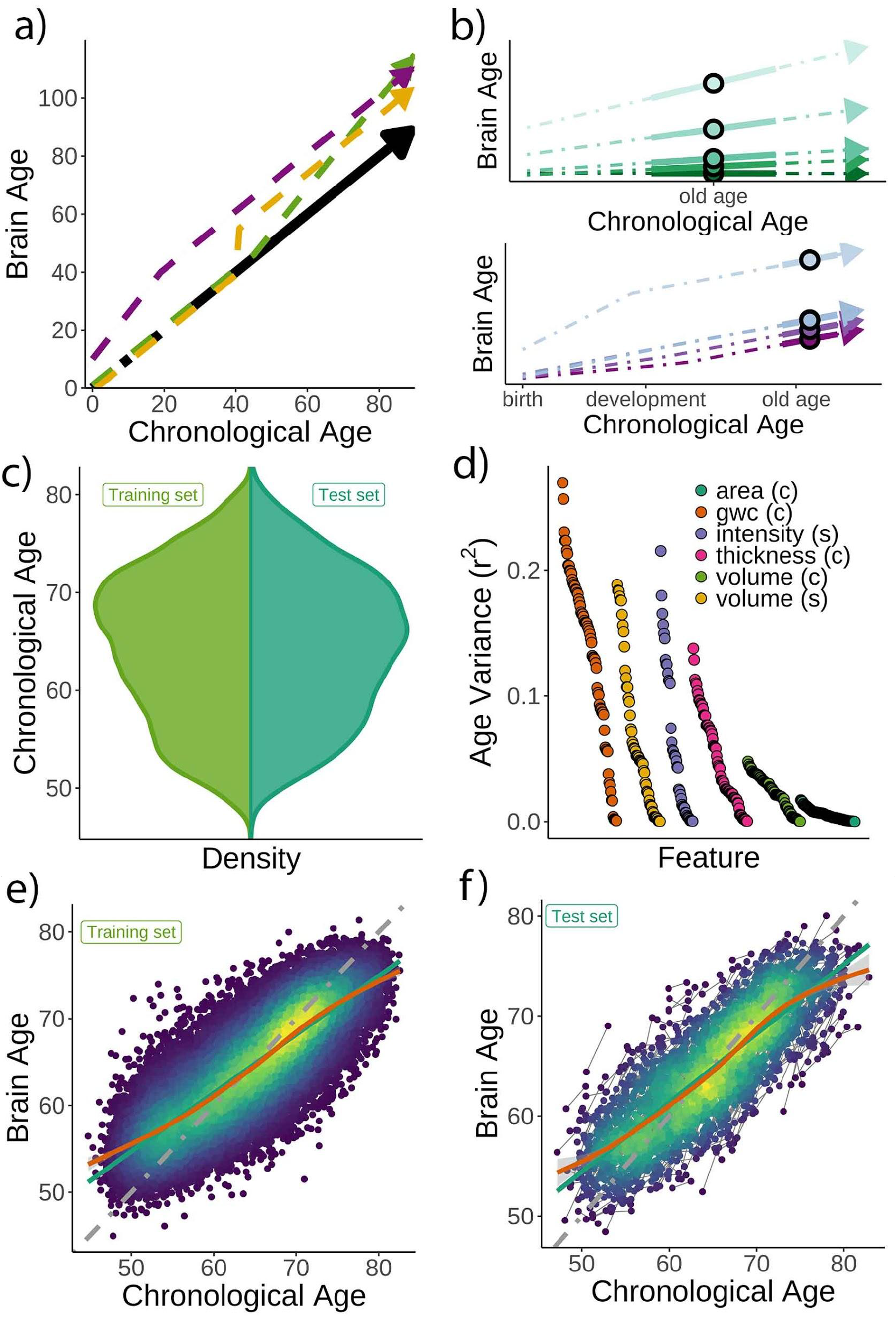
Theoretical expectations and study characteristics. **a)** Three hypothetical trajectories leading to higher *brain age delta*. Higher *brain age delta* can be explained by a steeper rate of neurobiological aging (green), distinct events that led to the accumulation of brain damage in the past (yellow), or early-life genetic and developmental factors (purple). The black arrow represents normative values of *brain age* through the lifespan. **b)** Brain aging (green) vs. early-life (blue-purple) accounts of *brain age* in older age. For the brain aging notion, cross-sectional *brain age* (points) relates to the slope of *brain age* as assessed by two or more observations across time (continuous line), reflecting ongoing differences in the rate of aging (dashed line, green scale). For the early-life notion, cross-sectional *brain age* (points), relates to early environmental, genetic, and/or developmental differences such as birth weight (blue-purple scale). **c)** Relative age distribution for the UK Biobank test and training datasets. **d)** Age variance explained (r^2^) for each MRI feature in the training dataset. Features are grouped by modality and ordered by the variance explained. **e)** *Brain age* model as estimated on the training (n = 38,682), and **f)** test datasets (participants = 1,372; two observations each). In **e)** and **f)**, lines represent the identity (grey), the linear (green), and the GAM (orange) fits of chronological age to *brain age*. Confidence intervals represent standard errors (SE). Note that plots show brain age prediction before age-bias correction. In **d)** gwc = gray-white matter contrast, (c) = cortical, and (s) = subcortical.

## Results

### Brain age prediction

Chronological age (**Fig. 1c**) was predicted based on regional and global features from structural T1-weighted (T1w) MRI, including cortical thickness, area, volume, and gray-white matter contrast, as well as subcortical volume and intensity imaging-derived phenotypes (|N| = 365). See list in **Supplementary Table 1, 2**, and **Fig. 1d** for pairwise correlations with age. The model was trained on 38,682 participants (age range = 44.8 - 82.6 years) with a single MRI from the UK Biobank^11^ dataset using gradient boosting as implemented in XGBoost (https://xgboost.readthedocs.io) and optimized using 10-fold cross-validation and a randomized hyper-parameters search. The trained model (**Fig. 1e**) was then used to predict *brain age* for an independent test dataset of 1,372 participants with two MRIs each (age range = 47.2 - 80.6 years, mean [SD] follow-up = 2.3 [0.1] years). The predictions revealed a high correlation between chronological and *brain age* (r = 0.82) with mean absolute error (MAE) = 3.31 years and root mean squared error (RMSE) = 4.14 years (**Fig. 1f**), comparable to other *brain age* models using UK Biobank MRI data^12^. We used generalized additive models (GAM) to correct for the brain-age bias, i.e., the *underestimation* of brain age in older individuals and vice versa^6^. *Brain age delta* was calculated as the residual from the GAM fit. *Brain age delta* at baseline and follow-up were strongly correlated (r = 0.81). To corroborate generalizability, we replicated our results using a different machine learning algorithm – a LASSO-based approach^12^ - and an independent longitudinal sample from the Lifebrain consortium^13^ with up to 11.2 years of follow-up (3,292 unique participants, age range = 18.0 - 94.4 years). See **Supplementary Fig. 1** and **Supplementary Table 3** for additional demographic information. All the code used to generate the results will be available at https://github.com/LCBC-UiO/VidalPineiro_BrainAge.

### Brain age does not strongly relate to the rate of brain aging

First, we tested whether cross-sectional *brain age delta* predicted *brain age delta*_*long*_ - i.e. annual rate of change in *brain age delta -* using linear models controlling for age, sex, scanning site, and estimated intracranial volume (eICV). We selected the centercept (*brain age delta* at mean chronological age), instead of baseline *brain age delta*, to avoid statistical dependency between indices. Cross-sectional and *brain age delta*_*long*_ were weakly, but negatively associated in the UK Biobank (β = -0.016 [± 0.008] *delta/*year, t (p) = -2.0 (.04), r^2^ = 0.002, **Fig. 2a**). Cross-sectional and *brain age delta*_*long*_ were unrelated using a LASSO regression approach (β = -0.003 [± 0.006] *delta/*year, t (p) = -0.5 (.65), r^2^ = .001, **Fig. 2b**), and in the Lifebrain replication sample (β = -0.007 [± 0.01] *delta/*year, t (p) = -0.6 (.53), r^2^ = 0.001, **Fig. 2c**). Post-hoc equivalence tests showed that positive relationships with β > 0.010 *delta/*year would be rejected in all three analyses thus confirming a lack of a meaningful relationship between cross-sectional and longitudinal *brain age* (**Methods** and **Supplementary Fig. 2**). UK Biobank (gradient boosting) results remained not significant when *brain age delta* was derived by timepoints 1 and 2 as two independent training sets (10-fold cross-validation; *uncorrected delta* values), thus avoiding potential confounds with age-bias correction (t (p) = 0.3 (.76)). Lifebrain results remained unaffected after including follow-up interval as an additional covariate or restricting the analysis to participants with long follow-up intervals (> 4 years) (n = 424). The relationship between cross-sectional and *brain age delta*_*long*_ was not significant in both cases (β = -0.008 [± 0.01] year/*delta*, t (p) = -0.7 (.45); β = -0.008 [± 0.007] *delta/*year, t (p) = -1.1 (.26)).

**Fig. 2.**
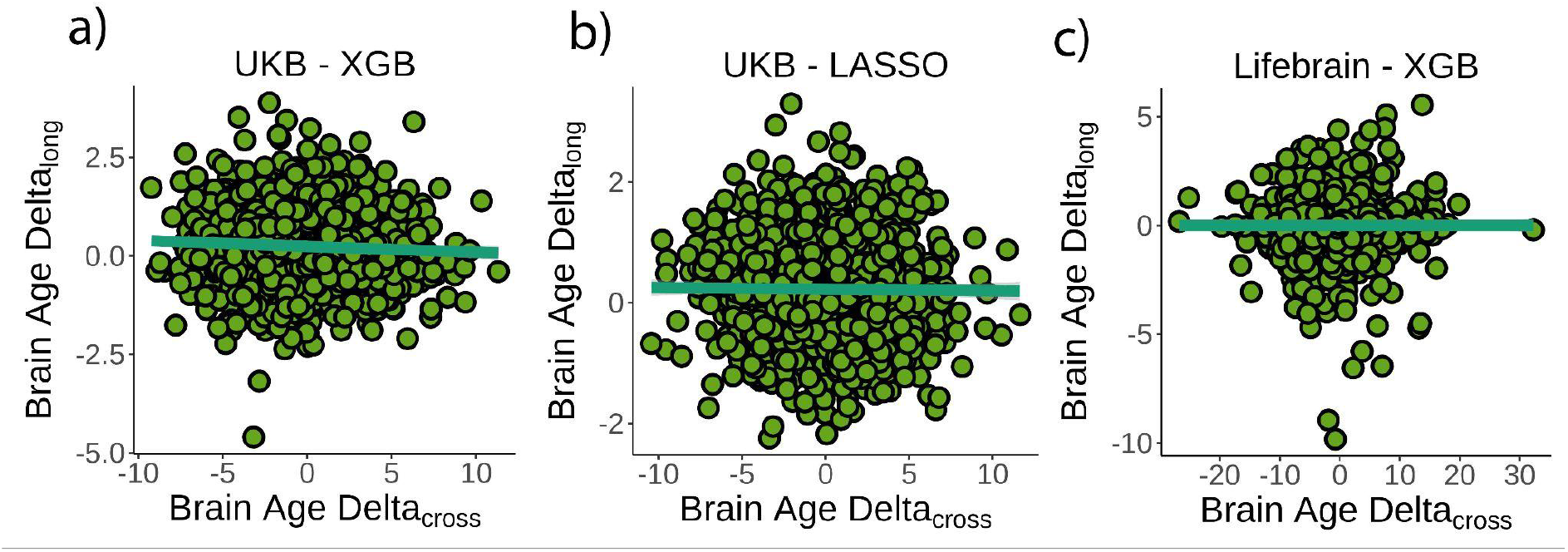
Relationship between cross-sectional and longitudinal *brain age delta*. **a)** Main analysis using the UK Biobank dataset and boosting gradient (n = 1372) (p = .04, r^2^ = 0.002). **b)** Replication analyses using a different training algorithm (LASSO; n = 1372) (p = .65, r^2^ = 0.001) and **c)** an independent dataset (Lifebrain; n = 1500) (p = 0.53, r^2^ = 0.001). XGB = boosting gradient as implemented in XGBoost. Confidence intervals represent SE. Longitudinal *brain age delta* (brain age delta_long_) refers to the rate of change in delta between baseline and follow-up MRI measurements. Cross-sectional *brain age delta* (brain age delta_cross_) refers to centercept *brain age delta*; i.e. at mean age.

We additionally tested whether cross-sectional and longitudinal *brain age delta (brain age delta*_*long*_) were associated with a composite measure of longitudinal brain change or with change in any of the structural MRI features. See **Methods** for details. Cross-sectional *brain age delta* was unrelated to a principal component of change (β = -0.009 [± 0.01] year, t (p) = -0.7 (.46), r^2^ = 0.001). We did not find a significant relationship when *brain age delta* was computed with neither a LASSO algorithm nor using the Lifebrain sample (β = -0.02 [± 0.01] year, t (p) = -1.7 (0.09), r^2^ = 0.002; β = 0.007 [± 0.006] year, t (p) = 1.3 (0.2), r^2^ = 0.001). In contrast, *brain age delta*_*long*_ was associated with a principal component of change in the UK Biobank dataset as well as in both replication analyses (all tests p < 0.001). See **Supplementary Fig. 3** for a visual representation. At a level of specific features, cross-sectional *brain age delta* was significantly related to change - in the expected direction - of features capturing lateral ventricle expansion and white matter hypointensities (p < 0.05 Bonferroni corrected). *Brain age delta*_*long*_ related to change in 45 of the features pertaining to four different modalities. The results were replicated both using the LASSO algorithm and the Lifebrain dataset (**Supplementary Fig. 4** and **Supplementary Table 4**).

Finally, we estimated the *rate of aging* effects using a cross-sectional model by estimating the scaling of the size of *delta* with age as defined in Smith and colleagues^6^. The scaling (λ) of *brain age delta* (**δ**) throughout the datasets’ age range was λ = 0.14 and 0.09 for the UK Biobank and the Lifebrain datasets. This corresponds to an increase in the spread of *brain age delta* of |**δ**|λ = .38 and .37 years - when moving from youngest to oldest - in the UK Biobank and the Lifebrain datasets suggesting that *brain age delta* only modestly reflects *rate of aging* effects.

### Brain age delta is associated with early life influences and polygenic scores for brain age

Next, we tested if birth weight was associated with *brain age delta* or change in *brain age delta*. Linear mixed models were used to fit time (from baseline; years), birth weight, and its interaction on *brain age delta*, using age at baseline, sex, scanning site, and eICV as covariates. Birth weight was significantly related to *brain age delta* (β = -0.70 [± 0.30] year/kg, t (p) = -2.3 (0.02), r^2^ = .009, **Fig. 3a**) but not to *delta* change (β = 0.02 [± .09] year/kg, t (p) = 0.3 (.79)). Birth weights were limited to normal variations at full-term (from 2.5 to 4.5 kg) (n = 770 unique individuals) but see **Supplementary Fig. 5** for results with varying cut-offs. The results were not affected by excluding individuals being part of multiple births (p = 0.02) and were replicated using the LASSO approach (**β** = -0.79 [± .29] year/kg, t (p) = -2.8 (0.006), r^2^ = 0.009, **Fig. 3b)**.

**Fig. 3.**
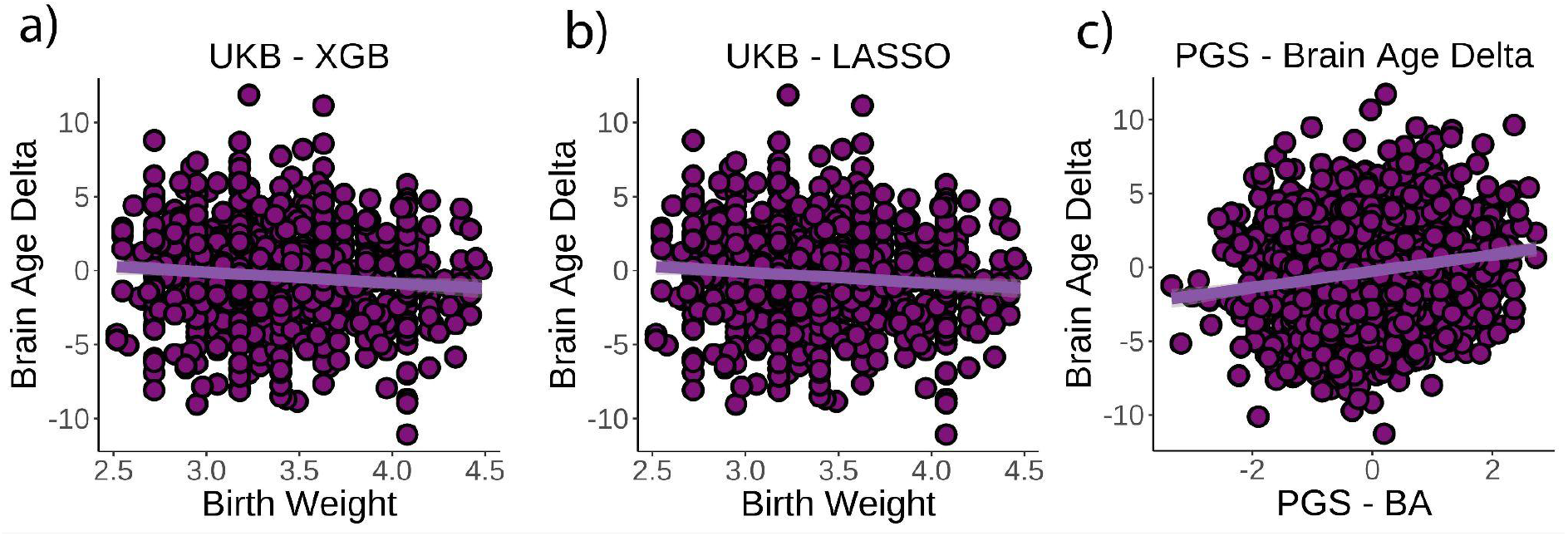
Relationship between cross-sectional *brain age delta and birth weight*. **a)** Main analysis using the UK Biobank dataset and boosting gradient (n = 770) (p = 0.02, r^2^ = 0.009). **b)** Replication analyses using a different training algorithm (LASSO) (n = 770) (p = 0.005, r^2^ = 0.009). **c)** Relationship between polygenic scores for *brain age delta* and *brain age delta* (n = 1,339) (p < 0.001, r^2^ = 0.02). XGB = boosting gradient as implemented in XGBoost. Confidence intervals represent SE.

Finally, we tested whether polygenic scores for *brain age delta* (PGS-BA) related to *brain age delta* and change in *brain age delta* (n = 1,339). PGS-BA was computed using a mixture-normal model based on a genome-wide association study (GWAS) of the *brain age delta* phenotype in the UK Biobank training dataset. To test the association, linear mixed models were used as above along with the top 10 genetic principal components to account for population structure. PGS-BA was positively associated with *brain age delta* (**β** = 0.54 [± 0.09] year, t (p) = 9.4 (< 0.001), r^2^ = 0.02, **Fig. 3c)** and negatively associated with *brain age delta* change (**β** = -0.06 [± 0.03] year, t (p) = -2.4 (0.02)) in the independent test dataset. Likewise, PGS-BA was associated with *brain age delta* derived from the LASSO algorithm (**β** = 0.53 [± 0.09] year, t (p) = 10.4 (< 0.001), r^2^ = 0.02) but not to *brain age delta* change (**β** = -0.001 [± .02] year, t (p) = 0.0 (1.0)). See **Supplementary Fig. 6** for GWAS association results. The association between PGS-BA and *brain age delta* remained significant when using as covariates the top 10 genetic components derived from the full UK Biobank sample (p < .001 in both analyses).

## Discussion

Altogether, these findings do not support the claim that the individual variations in the cross-sectional *brain age* metric captures across-subject differences in the ongoing rate of brain aging. Rather, *brain age* seems to reflect early-life influences, and only to a very modest degree reflects actual rate of brain change in middle and old adulthood. A lack of relationship between *brain age* and rate of brain aging can potentially be explained by the effect of circumscribed events such as isolated insults or detrimental lifestyles that occurred in the past resulting in higher, but not accelerating, *brain age*. Yet, variations in *brain age* can equally reflect developmental and early-life differences and show lifelong stability. *Brain-age* datasets and paradigms are generally ill-suited to disentangling these sources of variation but are often interpreted in line with the former. This assumes that variation in *brain age* largely results from the accumulation of damage and insults during the lifespan, with similar starting points for everyone. An exception is Elliott and colleagues^3^, who found that middle-aged individuals with higher brain age already exhibited poorer cognitive function and brain health at age three years. This fits a robust corpus of literature showing effects of lifelong, stable influences as indexed by childhood IQ^14^, genetics^10^, and neonatal characteristics^8^ on brain and cognitive variation in old age.

Strictly speaking, *brain age delta* is a prediction error from a model that maximizes the prediction of age in cross-sectional data. Prediction errors also reflect noise, attenuating any relation between cross-sectional and longitudinal *brain age*. Given that delta_long_ is estimated as the difference between two delta_cross_ estimates, it will hence have higher noise than the cross-sectional estimates reducing the power in identifying potential associations between longitudinal and cross-sectional delta; note also the relatively short interscan interval in UK Biobank (≈2y). However, replication (of our null results) in the Lifebrain sample with more observations and longer follow-up reduces the likelihood of noise as the main factor behind the lack of relationship. Furthermore, previous studies have found that changes in *brain age* are partly heritable^15^ and relate to for instance cardiometabolic risk factors^16^, suggesting that it captures biologically relevant signals, although with substantially different origins from cross-sectional *brain age*. It has been argued that at a population level, brain age captures modest *rate of aging* effects as *brain age delta* spreads with increasing age^6^. Here, we found a similar degree of *delta* spreading in our *brain age* metrics. Likewise, our secondary analyses suggested *brain age* related to change in few specific neuroimaging features, i.e. ventricular expansion and white matter hypointensities, though not to any composite score. Thus, both results are compatible and converge towards *brain age* as a real but relatively modest metric for capturing ongoing brain change. The largest part of interindividual variation in *brain age delta*, instead, largely originates before the sample lower bound (⪝ 18 and 45 years for the Lifebrain and UK Biobank datasets). Other multivariate approaches might be better equipped for capturing the dynamics of the aging brain. Using independent component analysis, a recent study found that - compared to a single *brain age* score - distinct modes of multimodal brain variation better reflect both the genetic make-up and ongoing aging effects, with a subset of *modes* showing significant spreading of *delta* with age^5^.

The degree to which *brain age* reflects ongoing effects likely depends on the specific features, modalities, and algorithms employed and is constrained by model properties such as prediction accuracy and homoscedasticity. Yet, without longitudinal imaging, one should not interpret brain age as accelerated aging. Our results align with theoretical claims and empirical observations that covariance structures capturing differences between individuals do not necessarily generalize to covariance structures within individuals^17,18^. Also, associations of *brain age* with other bodily markers of aging or with cognitive decline have yielded mixed support for cross-sectional *brain age* as a marker of individual differences in brain aging^2,3,19^. Strong relationships between cross-sectional and longitudinal *brain age* may thus be restricted to specific disease groups such as Alzheimer’s disease patients^19^ where interindividual brain variation is dominated by the prevailing loss of brain structural integrity.

The results further showed that birth weight, which reflects differences in genetic propensities and prenatal environment^20^, explained a modest portion of the variance in *brain age*. Subtle variations in birth weight are associated with brain structure early in life and present throughout the lifespan^8^. This association should be considered as *proof-of-concept* that the metric of *brain age* reflects the distant past more than presently ongoing events in the morphological structure of the brain. This was confirmed by the consistent association between PGS-BA and *brain age delta* but not with *brain age delta* change. Since PGS-BA was computed based on cross-sectional *brain age delta*, this relationship may not be surprising, but still suggests a different genetic foundation for longitudinal *brain age*. These findings link with evidence that brain development is strongly influenced by genetic architecture that, in interaction with environmental factors, lead to substantial, long-lasting effects on brain structure. By contrast, aging mechanisms seem to be more related to limitations of maintenance and repair functions and have a more stochastic nature^21^.

As time from birth increases, chronological age as a marker of individual development is reduced. The results call for caution in interpreting brain-derived indices of aging based on cross-sectional MRI data and underscores the need to rely on longitudinal data whenever the goal is to understand the trajectories of brain and cognition in aging.

## Methods

### Participants and Samples

The main sample was drawn from the UK Biobank neuroimaging branch (https://www.ukbiobank.ac.uk/)^11^. 38,682 individuals had MRI available at a single time point and were used as the training dataset. 1,372 individuals had longitudinal data and were used as the test dataset. The present analyses were conducted under data application number 32,048. The Lifebrain dataset^13^ included datasets from 5 different major European Lifespan cohorts: the Center for Lifespan Changes in Brain and Cognition cohort (LCBC, Oslo)^8^, the Cambridge Center for Aging and Neuroscience study (Cam-CAN)^22,23^, the Berlin Study of Aging-II (Base-II)^24^, the University of Barcelona cohort (UB)^25,26^, and the BETULA project (Umeå)^27^. Furthermore, we included data from the Australian Imaging Biomarkers and Lifestyle flagship study of ageing (AIBL)^28^. In addition to cohort-specific inclusion and exclusion criteria, individuals aged < 18 years, or with evidence of mild cognitive impairment, or Alzheimer’s Disease were excluded from the analyses. 1,792 individuals with only one available scan were used for the Lifebrain training dataset. 1,500 individuals with available follow-up of > 0.4 years were included in the test dataset. Individuals had between 2 and 8 available scans each. Sample demographics for the UK Biobank and the Lifebrain samples are provided in **Supplementary Table 3**. See also **Fig. 1c** and **Supplementary Fig. 1** for a visual representation of the age distribution in the UK Biobank and the Lifebrain datasets. UK Biobank (North West Multi-Center Research Ethics Commitee [MREC]; see also https://www.ukbiobank.ac.uk/the-ethics-and-governance-council) and the different cohorts of the Lifebrain replication dataset (**Supplementary Table 5**) have ethical approval from the respective regional ethics committees. All participants provided informed consent.

### MRI acquisition and preprocessing

See https://biobank.ctsu.ox.ac.uk/crystal/crystal/docs/brain_mri.pdf for details on the UK Biobank T1-weighted (T1w) MRI acquisition. UK Biobank and Lifebrain MRI data were acquired with 3 and 10 different scanners, respectively. T1w MRI acquisition parameters for both the Lifebrain and the UK Biobank are summarized in **Supplementary Table 6**.

We used summary regional and global metrics derived from T1w data. For UK Biobank we used the imaging-derived phenotypes developed centrally by UK Biobank researchers^11^ and distributed via the data showcase (http://biobank.ctsu.ox.ac.uk/crystal/index.cgi). See preprocessing details in https://biobank.ctsu.ox.ac.uk/crystal/crystal/docs/brain_mri.pdf. This procedure yielded 365 structural MRI features, partitioned in 68 features of cortical thickness, area, and gray-white matter contrast, 66 features of cortical volume, 41 features of subcortical intensity, and 54 features of subcortical volume. See the list of features in **Supplementary Table 1** and **2**. Lifebrain data were processed on the Colossus processing cluster, University of Oslo. Similar to the UK Biobank pipeline, we used the fully automated longitudinal FreeSurfer v.6.0. pipeline^29^ for cortical reconstruction and subcortical segmentation of the structural T1w data (http://surfer.nmr.mgh.harvard.edu/fswiki)^30–32^ and used similar atlases for structural segmentation and feature extraction.

### Birth weight

We used birth weight (kg) from the UK Biobank (*field #20022*). Participants were asked to enter their birth weight at the initial assessment visit, the first repeat assessment visit, or the first imaging visit. In the case of multiple birth weight instances, we used the latest available input. n = 894 participants from the test dataset had available data on birth weight. The main analysis was constrained to normal variations in birth weight between 2.5 and 4.5 kg (n = 770)^33^ due to lower reliability of extreme scores and to tentatively remove participants with severe medical complications associated with prematurity.

### Genetic preprocessing

Detailed information on genotyping, imputation, and quality control was published by Bycroft and colleagues^34^. For genetic analyses, we only included participants with both genotypes and MRI scans. Following the recommendations from the UK Biobank website, we excluded individuals with failed genotyping, that had abnormal heterozygosity status, or that withdrew their consents. We also removed participants that were genetically related – up to the third degree – to at least another participant as estimated by the kinship coefficients as implemented in PLINK^35^. For the genome-wide association study (GWAS) we used 38,163 individuals from the training dataset. Polygenic risk scores were computed using the test dataset consisting of 1,339 individuals with longitudinal MRI.

#### GWAS

We performed GWAS analysis on the training dataset and the *brain age delta-semi-*corrected phenotype using the imputed UK Biobank genotypes. To control for subtle effects of population stratification in the dataset, we computed the top 10 principal components (PCs) using the PLINK command *–pca* on a decorrelated set of autosome single nucleotide polymorphisms (SNPs). The set of SNPs (n=101,797) were generated by using the PLINK command, *--maf 0*.*05, --hwe 1e*^*-6*^, *--indep-pairwise 100 50 0*.*1*. The *–glm* function from PLINK was used to perform GWAS on about 9 million autosomal SNPs, including age, sex, and the top 10 PCs as covariates. See Manhattan and quantile-quantile (QQ) plots in **Supplementary Fig. 6**. Note that our results corroborated the same association region reported in Jonsson and colleagues^36^ with a smaller sample.

#### Polygenic scores (PGS)

The GWAS results for the training dataset were used to compute PGS (PGS-BA) in the independent test dataset (n = 1,339 participants). We used the recently developed method PRS-CS^37^ to estimate the posterior effect sizes of SNPs that were shown to have high quality in the HapMap data^38^. Rather than estimating the polygenicity of *brain age delta* from our data, we assumed a highly polygenic architecture for *brain age delta* by setting the parameter *--phi=0*.*01*^39^. *The remaining parameters of PRS-CS were set to the default values. PGS was based on 654,725 SNPs and was computed on the independent test data using the --score* function from PLINK. SNPs were aligned with HapMap 3 SNPs (autosome only as provided by PRC-CS) and posterior effects were estimated. We also computed the population structures PCs’ in the test dataset using the same procedure as in the training dataset.

### Statistical analyses

All statistical analyses were run with R version 3.6.3 https://www.r-project.org/. We used the UK Biobank as the main sample and the Lifebrain cohort for independent replication. The main description refers to the UK Biobank pipeline, though Lifebrain replication followed identical steps unless otherwise stated. For replication across machine learning pipelines, we used a LASSO regression approach for age prediction, adapted from https://james-cole.github.io/UKBiobank-Brain-Age/. See more details in Cole, 2020^12^. The correlation between LASSO-based and Gradient Boosting-based *brain age deltas* was .80.

#### Brain age prediction

We used machine learning to estimate each individuals’ *brain age* based on a set of regional and global features extracted from T1w sequences. We estimated *brain age* using gradient tree boosting (https://xgboost.readthedocs.io). We used participants with only one MRI scan for the training dataset (n = 36,682) and participants with longitudinal data as test dataset (n = 1,372). All variables were scaled prior to any analyses using the training dataset metrics as reference.

The model was optimized in the training set using a 10-fold cross-validation randomized hyper-parameters search (50 iterations). The hyper-parameters explored were number of estimators [100, 600, 50], learning rate (0.01, 0.05, 0.1, 0.15, 0.2), maximum depth [2, 8, 1], gamma regularisation parameter [0.5, 1.5, 0.5], and min child weight [1, 4, 1]. The remaining parameters were left to default. The optimal parameters were: number of estimators = 500, learning rate = 0.1, maximum depth = 5, gamma = 1, and min child weight = 4 predicting r^2^ = 0.68 variance in chronological age with mean absolute error (MAE) = 3.41 and root mean squared error (RMSE) = 4.29. See visual representation in **Fig. 1f**.

Next, we recomputed the machine learning model using the entire training dataset and the optimal hyper-parameters and used it to predict *brain age* for the test dataset (**Fig. 1e)**. These metrics are similar or better than other *brain age* models using UK Biobank MRI data^12,40^ and the cross-validation diagnostics. We used GAM to correct for the brain-age bias estimation^6^; r = -0.54 for the test dataset. Note that we used GAM fittings as estimated in the training dataset so *delta* values in the test dataset are not centered to 0. *Brain age delta* was estimated as the GAM residual. The correlation between *brain age delta* corrected based on the training vs. the test fit was r > 0.99. Also, GAM-based bias correction led to similar brain age delta estimations to linear and quadratic-based corrections (r > 0.99). The diagnostics for LASSO-based model were as follows: variance explained (r^2^)= 0.83 / 0.83; MAE = 3.36 / 3.28; RMSE = 4.21 / 4.04; age-bias = -.56 / -.52 for the training and predicted datasets. See representation of the *brain age* prediction in **Supplementary Fig. 7**.

#### Higher level analysis

##### Relationship between cross-sectional and longitudinal brain age

For each participant, we computed the mean *brain age delta* across the two MRI time points and the yearly rate of change (*brain age delta*_*long*_). We selected mean, instead of baseline *brain age delta*, to avoid statistical dependency between both indices^41,42^. *Brain age delta*_*long*_ was fitted by mean *brain age delta* using a linear regression model, which accounted for age, sex, site, and eICV. We used mean eICV across both time points.

##### Relationship between brain age delta and change in brain features

For each participant, we computed the yearly rate of change in all the *raw* neuroimaging features and tested whether change was significantly different from 0 (one-sample t-test, p < 0.05, Bonferroni-corrected) **(Supplementary Fig. 4, Supplementary Table 4)**. Features with significant change over time were fed into a principal component analysis (uncentered). The first component, explaining ⋍20% of the variance both in the UK Biobank and the Lifebrain datasets, was selected for further analysis. Although it did not qualitatively affect the results, we removed two and three extreme outliers from the UK Biobank and Lifebrain datasets (score > 10). See **Supplementary Table 4** for component weights. Finally, we tested whether cross-sectional and *brain age delta*_*long*_ predicted brain change as quantified both by the first component analysis and change in each of the *raw* neuroimaging features (p < 0.05, Bonferroni-corrected) using the same models described above.

##### Spreading of brain age delta with age

Further, we estimated the degree to which *brain age delta* reflects *rate of aging* using a cross-sectional model proposed by Smith^6^ which estimates the scaling of *brain age delta* through the datasets’ age range. The scaling is estimated by λ in **δ** = **δ**_0_(1 +λY_0_) where **δ** is *brain age delta*, Y_0_ is a linear mapping of chronological age into the range 0:1, and |**δ**_0_| relates to *brain age delta* distribution in the youngest participants. The *spread* of *brain age delta* throughout the datasets’ age range can then be expressed as |**δ**_0_|λ (years).

##### Relationship between brain age PGS and cross-sectional and longitudinal brain age

This association was tested using linear mixed models with time from baseline (years), PGS-BA, and its interaction on *brain age delta*. Age at baseline, sex, site, eICV, and the 10 first principal components for population structure were used as covariates. The principal components of population structure were added to minimize false positives associated with any form of relatedness within the sample. *Effects of birth weight on brain age*. Linear mixed models were used to fit time, birth weight, and its interaction on *brain age delta*, using age at baseline, sex, site, and eICV as covariates. We explored the consistency of the results by modifying the birth weight limits in a grid-like fashion [0.5, 2.7, 0.025] and [4.2, 6.5, 0.025] for minimum and maximum birth weight (**Supplementary Fig. 5**). Self-reported birth weight is a reliable estimate of actual birth weight. However, extreme values are either misestimated or reflect profound gestational abnormalities^43,44^.

##### Equivalence tests

Post-hoc equivalence tests were carried to test for the absence of a relationship between cross-sectional and *brain age delta* _*long*_ ^45^. *Specifically, we used inferiority tests, to test whether a null hypothesis of an effect as least as large as Δ (in years/delta*) could be rejected. We re-run the three main models assessing a relationship between cross-sectional and longitudinal *brain age delta* (UK Biobank trained with boosting gradient, UK Biobank trained with LASSO, and Lifebrain trained with boosting gradient) varying the right-hand-side test (Δ) [-0.02, 0.05, 0.001] (p < 0.05, one-tailed) (**Supplementary Fig. 2**).

Assumptions were checked for the main statistical tests using plot diagnostics. Variance explained for single terms refers to unique variance (UVE), which is defined as the difference in explained variance between the full model and the model without the term of interest. For linear mixed models, UVE was estimated as implemented in the *MuMIn* r-package.

#### Lifebrain-specific steps

<u>*Features*</u>. The Lifebrain cohort included |N| = 372 features. It included 8 new features compared to the UK Biobank dataset, whereas one feature was excluded (new features: left and right temporal pole area volume and thickness, cerebral white matter volume, cortex volume; excluded feature: ventricle choroid). See age-variance explained in each feature in **Supplementary Table 1** and **2** as estimated with GAMs. <u>*Quality control*</u>. Prior to any analysis, we tentatively removed observations for which > 5% of the features fell above or below 5 SD from the sample mean. The application of this arbitrary high threshold led to the removal of 10 observations. We considered these MRI data to be extreme outliers and likely to be artifactual and/or contaminated by important sources of noise. Also, before brain prediction, we tentatively removed variance associated with the different scanners using generalized additive mixed models (GAMM) and controlling for age as a smooth factor and a subject-identifier as random intercept. This correction was performed due to differences in age distribution by scanner and lack of across scanner calibration. <u>*Hyperparameter search and model diagnostics*</u>. The optimal parameters for the Lifebrain replication sample were: number of estimators = 600, learning rate = 0.05, maximum depth = 4, gamma = 1.5, and min child weight = 1. Using cross-validation, the model predicted r^2^ = 0.92 of the age-variance with MAE = 4.75 and RMSE = 6.31. Brain age was underestimated in older age (bias r = -0.33). <u>*Model prediction*</u>. The age-variance explained by *brain age* was r = 0.90 with MAE = 4.68 and RMSE = 6.06. *Brain age* was underestimated in older age (bias r = -0.25) (**Supplementary Fig. 7**). <u>*Higher level-analysis*</u>. For each individual, mean *brain age delta* was considered as the grand-mean *brain age delta* across the different MRI time points. To compute *brain age delta*_*long*_ we set for each participant a linear regression model with observations equal to the number of time points that fitted *brain age delta* by time since the initial visit. Slope indexed change in *brain age delta*/year. The relationship between mean and *brain age delta*_*long*_ was tested using linear mixed models controlling for age, sex, and eICV as fixed effects, and using a site identifier as a random intercept. Likewise, linear mixed models were used to test the relationship between *brain age delta* and change in brain features. Note that Eicv was identical across timepoints as a result of being estimated through the longitudinal FreeSurfer pipeline. We could not obtain the required information on genetics and birth weight to replicate the analyses supporting the *early-life* account.

## Data availability

The raw data were gathered from the UK Biobank, the Lifebrain cohort, and the AIBL. Raw data requests are specific to each cohort. UK Biobank and AIBL data are available upon application to UK Biobank and at https://aibl.csiro.au upon corresponding approvals. For the Lifebrain cohorts, requests for raw MRI data should be submitted to the corresponding principal investigator. See contact details in **Supplementary Table 5**. Note that MRI data availability for some individuals may be restricted as participants did not consent to share publicly their data. Different restrictions and sample agreements might be required.

## Code availability

Statistical analyses in this manuscript will be available at https://github.com/LCBC-UiO/VidalPineiro_BrainAge. All analyses were performed in R 3.6.3. The scripts were run on the Colossus processing cluster, University of Oslo. UK Biobanks’ data acquisition, MRI preprocessing, and feature generation pipelines are freely available (https://www.fmrib.ox.ac.uk/ukbiobank). For the Lifebrain cohorts, the image acquisition details are summarized in **Supplementary Table 6**. MRI preprocessing and feature generation scripts were performed with the freely available FreeSurfer software (https://surfer.nmr.mgh.harvard.edu/). For bash-sourcing scripts, please contact the corresponding author.

## Acknowledgments

This work was supported by the Lifebrain project, funded by the EU Horizon 2020 Grant: ‘Healthy minds 0–100 years: Optimising the use of European brain imaging cohorts (‘Lifebrain’).” Grant agreement number: 732592 (to K.B.W.). In addition, the different sub-studies are supported by different sources. LCBC: The European Research Council under grant agreements 283634, 725025 (to A.M.F.) and 313440 (to K.B.W.), as well as the Norwegian Research Council (to A.M.F., K.B.W.), the National Association for Public Health’s dementia research program (A.M.F.) and the Peder Sather foundation (to D.V.P.). Betula: a scholar grant from the Knut and Alice Wallenberg (KAW) foundation to L.N. Barcelona: D.B.F. was funded by an ICREA Academia Award. D.B.F, B.S., and C.J. acknowledge the CERCA Programme/Generalitat de Catalunya and are supported by María de Maeztu Unit of Excellence (Institute of Neurosciences, University of Barcelona) MDM-2017-0729, Ministry of Science, Innovation and Universities. BASE-II has been supported by the German Federal Ministry of Education and Research under grant numbers 16SV5537/16SV5837/16SV5538/16SV5536K/01UW0808/01UW0706/01GL1716A/01GL1716B, and S.K. has received support from the European Research Council under grant agreement 677804. The Wellcome Centre for Integrative Neuroimaging is supported by core funding from award 203139/Z/16/Z from the Wellcome Trust. Drs Suri and Zsoldos were funded by an award the UK Medical Research Council (G1001354) and the HDH Wills 1965 Charitable Trust (1117747). Dr Suri is now funded by the UK Alzheimer’s Society Research Fellowship (Grant Ref 441); Suri is supported by the NIHR Oxford Health Biomedical Research Centre. Data used in the preparation of this article were partially obtained from the AIBL funded by the Commonwealth Scientific and Industrial Research Organisation (CSIRO), which was made available at the ADNI database (www.loni.usc.edu/ADNI). UK Biobank is generously supported by its founding funders the Wellcome Trust and UK Medical Research Council, as well as the Department of Health, Scottish Government, the Northwest Regional Development Agency, British Heart Foundation and Cancer Research UK. The organisation has over 150 dedicated members of staff, based in multiple locations across the UK.

## Author contributions

D.V.P, A.M.F, L.N., and K.B.W. designed and conceived the study; AIBL, B.S., C.J., S.S., S.D., R.W. and S.K.K. collected data; D.V.P, A.M.M., I.K.A., E.L., F.M., J.M.R, Y.W., L.B., and Ø.S. performed the analyses and created the figures; D.V.P, A.M.F, K.B.W., L.N., D.B.F, K.S.K., C.A.D., S.K., A.M.B., W.F.C.B., U.L., K.P.E., R.H., R.A.K., E.Zs., S.M.S, A.Z., and Y.W. interpreted the results. The paper was written by D.V.P. and A.M.F with substantial input from all the authors.

## References

1. Cole, J.H. & Franke, K. Trends Neurosci 40, 681–690 (2017).

2. Cole, J.H. et al. Molecular Psychiatry 23, 1385–1392 (2018).

3. Elliott, M.L. et al. Mol Psychiatry (2019).doi:10.1038/s41380-019-0626-7

4. Franke, K. & Gaser, C. Front. Neurol. 10, (2019).

5. Smith, S.M. et al. Elife 9, (2020).

6. Smith, S.M., Vidaurre, D., Alfaro-Almagro, F., Nichols, T.E. & Miller, K.L. NeuroImage 200, 528–539 (2019).

7. Deary, I.J. Gerontology 58, 545–553 (2012).

8. Walhovd, K.B. et al. Proc. Natl. Acad. Sci. U.S.A. 113, 9357–9362 (2016).

9. Walhovd, K.B. et al. Proc Natl Acad Sci U S A 109, 20089–20094 (2012).

10. Walhovd, K.B. et al. Neurology Genetics 6, (2020).

11. Miller, K.L. et al. Nat. Neurosci. 19, 1523–1536 (2016).

12. Cole, J.H. Neurobiology of Aging 92, 34–42 (2020).

13. Walhovd, K.B. et al. Eur. Psychiatry 50, 47–56 (2018).

14. Karama, S. et al. Mol. Psychiatry 19, 555–559 (2014).

15. Brouwer, R.M. et al. Cereb Cortex doi:10.1093/cercor/bhaa296

16. Beck, D. et al. medRxiv 2021.02.25.21252272 (2021).doi:10.1101/2021.02.25.21252272

17. Molenaar, P.C.M. Measurement: Interdisciplinary Research and Perspectives 2, 201–218 (2004).

18. Schmiedek, F., Lövdén, M., von Oertzen, T. & Lindenberger, U. PeerJ 8, e9290 (2020).

19. Franke, K. & Gaser, C. GeroPsych: The Journal of Gerontopsychology and Geriatric Psychiatry 25, 235–245 (2012).

20. Gielen, M. et al. Behav Genet 38, 44–54 (2008).

21. Kirkwood, T.B.L. Cell 120, 437–447 (2005).

22. Shafto, M.A. et al. BMC Neurol 14, (2014).

23. Taylor, J.R. et al. NeuroImage 144, 262–269 (2017).

24. Bertram, L. et al. Int J Epidemiol 43, 703–712 (2014).

25. Rajaram, S. et al. Front Aging Neurosci 8, (2017).

26. Vidal-Piñeiro, D. et al. Brain Stimul 7, 287–296 (2014).

27. Nilsson, L.-G. et al. Aging, Neuropsychology, and Cognition 11, 134–148 (2004).

28. Ellis, K.A. et al. Int Psychogeriatr 21, 672–687 (2009).

29. Reuter, M., Schmansky, N.J., Rosas, H.D. & Fischl, B. NeuroImage 61, 1402–1418 (2012).

30. Dale, A.M., Fischl, B. & Sereno, M.I. Neuroimage 9, 179–194 (1999).

31. Fischl, B., Sereno, M.I. & Dale, A.M. Neuroimage 9, 195–207 (1999).

32. Fischl, B. & Dale, A.M. Proc. Natl. Acad. Sci. U.S.A. 97, 11050–11055 (2000).

33. Walhovd, K.B. et al. Proc. Natl. Acad. Sci. U.S.A. 109, 20089–20094 (2012).

34. Bycroft, C. et al. Nature 562, 203–209 (2018).

35. Chang, C.C. et al. Gigascience 4, 7 (2015).

36. Jonsson, B.A. et al. Nature Communications 10, 5409 (2019).

37. Ge, T., Chen, C.-Y., Ni, Y., Feng, Y.-C.A. & Smoller, J.W. Nature Communications 10, 1776 (2019).

38. International HapMap 3 Consortium et al. Nature 467, 52–58 (2010).

39. Boyle, E.A., Li, Y.I. & Pritchard, J.K. Cell 169, 1177–1186 (2017).

40. Lange, A.-M.G. de et al. PNAS 116, 22341–22346 (2019).

41. Rogosa, D.R. & Willett, J.B. Psychometrika 50, 203–228 (1985).

42. Wainer, H. Psychol Sci 11, 434–436 (2000).

43. Nilsen, T.S., Kutschke, J., Brandt, I. & Harris, J.R. Twin Res Hum Genet 20, 406–413 (2017).

44. Tehranifar, P., Liao, Y., Flom, J.D. & Terry, M.B. Am J Epidemiol 170, 910–917 (2009).

45. Lakens, D., Scheel, A.M. & Isager, P.M. Advances in Methods and Practices in Psychological Science 1, 259–269 (2018).

