## Supplementary Information for "Individual variations in “Brain age” relate to early life factors more than to longitudinal brain change"

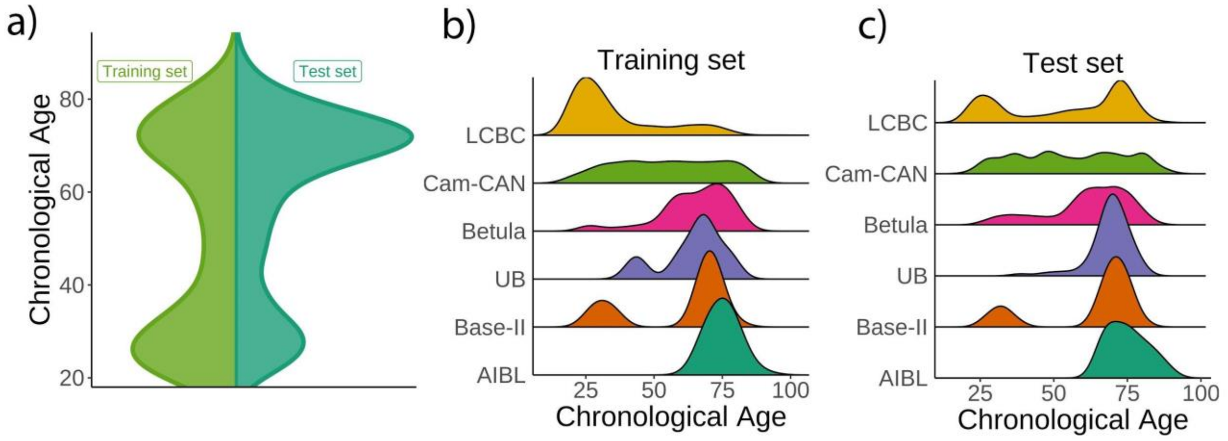

**Supplementary Fig. 1. Age distribution for the Lifebrain replication dataset. a)** Relative age distribution for the Lifebrain training and test datasets. Relative age distribution for the different cohorts of the Lifebrain **b)** training and **c)** test datasets.

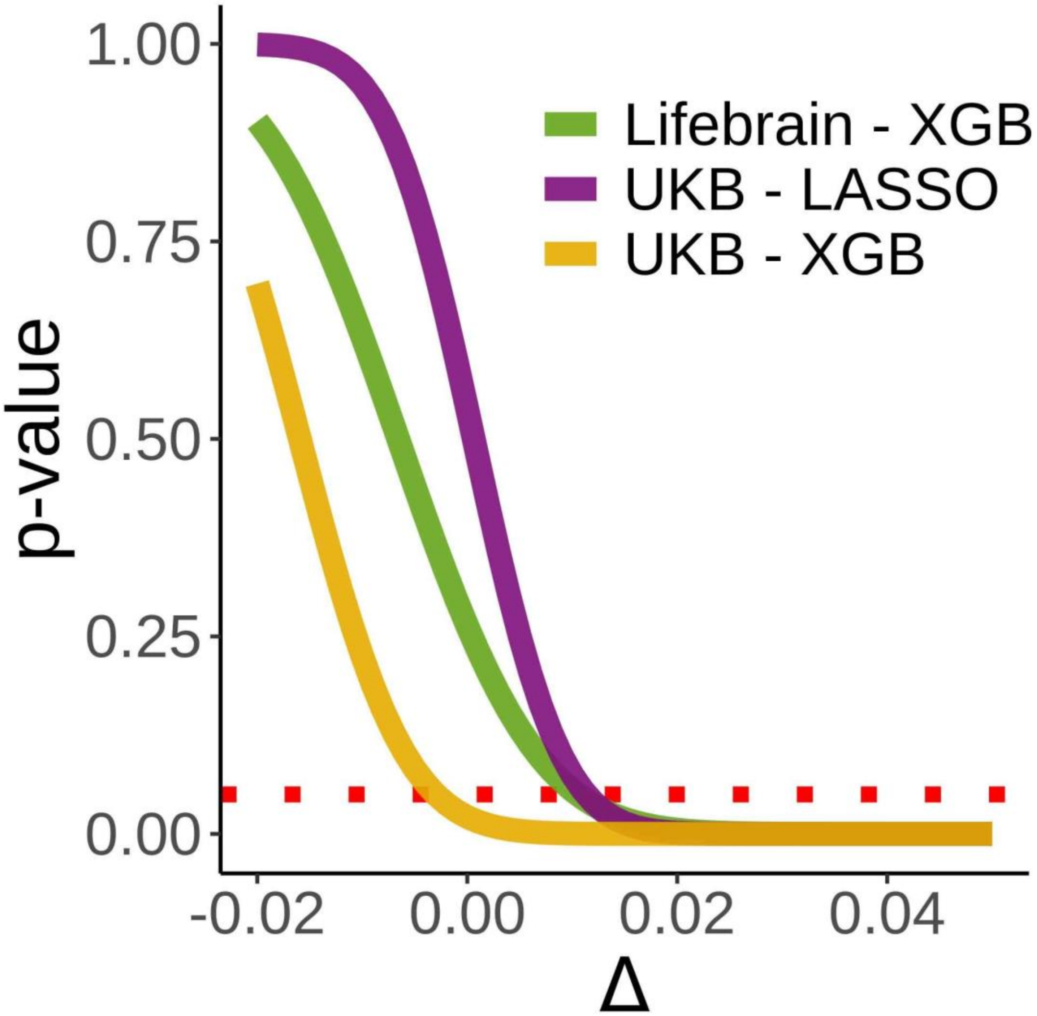

**Supplementary Fig. 2. Equivalence tests.** Inferiority tests for the three main models used to assess the relationship between cross-sectional and *brain age delta*$_{long}$. Inferiority tests test whether a null hypothesis of an effect as large as Δ can be rejected. In the x-axis, Δ reflects the null hypothesis as βetas (years/*delta*). A null hypothesis of an effect at least as large as 0.11 years/*delta* can be rejected ($p < 0.05$) in all three tests. Δ has been evaluated at [-0.02, 0.05, 0.001]. The dashed red line indicates a $p = 0.05$ criterion for the null hypothesis rejection.

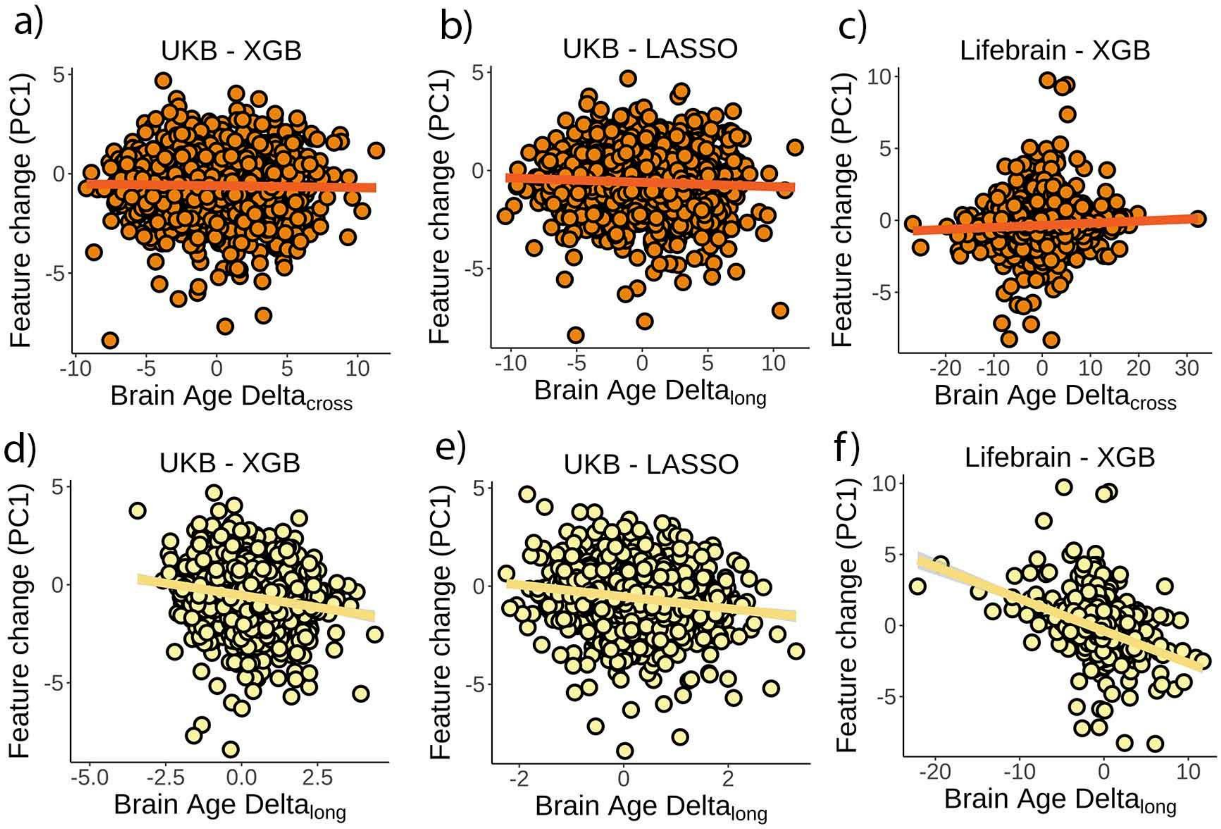

**Supplementary Fig. 3. Relationship between *brain age delta* and composite measures of change.** Relationship between a composite measure of change as capture by the first principal component on feature change and cross-sectional *brain age delta* in a) the UK Biobank and boosting gradient, b) the UK Biobank and the LASSO algorithm, and c) the Lifebrain dataset. Relationship between the composite measure of change and (longitudinal) *brain age delta$_{long}$* in d) the UK Biobank and boosting gradient, e) the UK Biobank and the LASSO algorithm, and f) the Lifebrain dataset. Negative values in the principal component reflect brain decline (e.g. steeper cortical thinning, higher ventricle volume, etc.). n = 1,369 and 1,497 for the UK Biobank and the Lifebrain datasets.

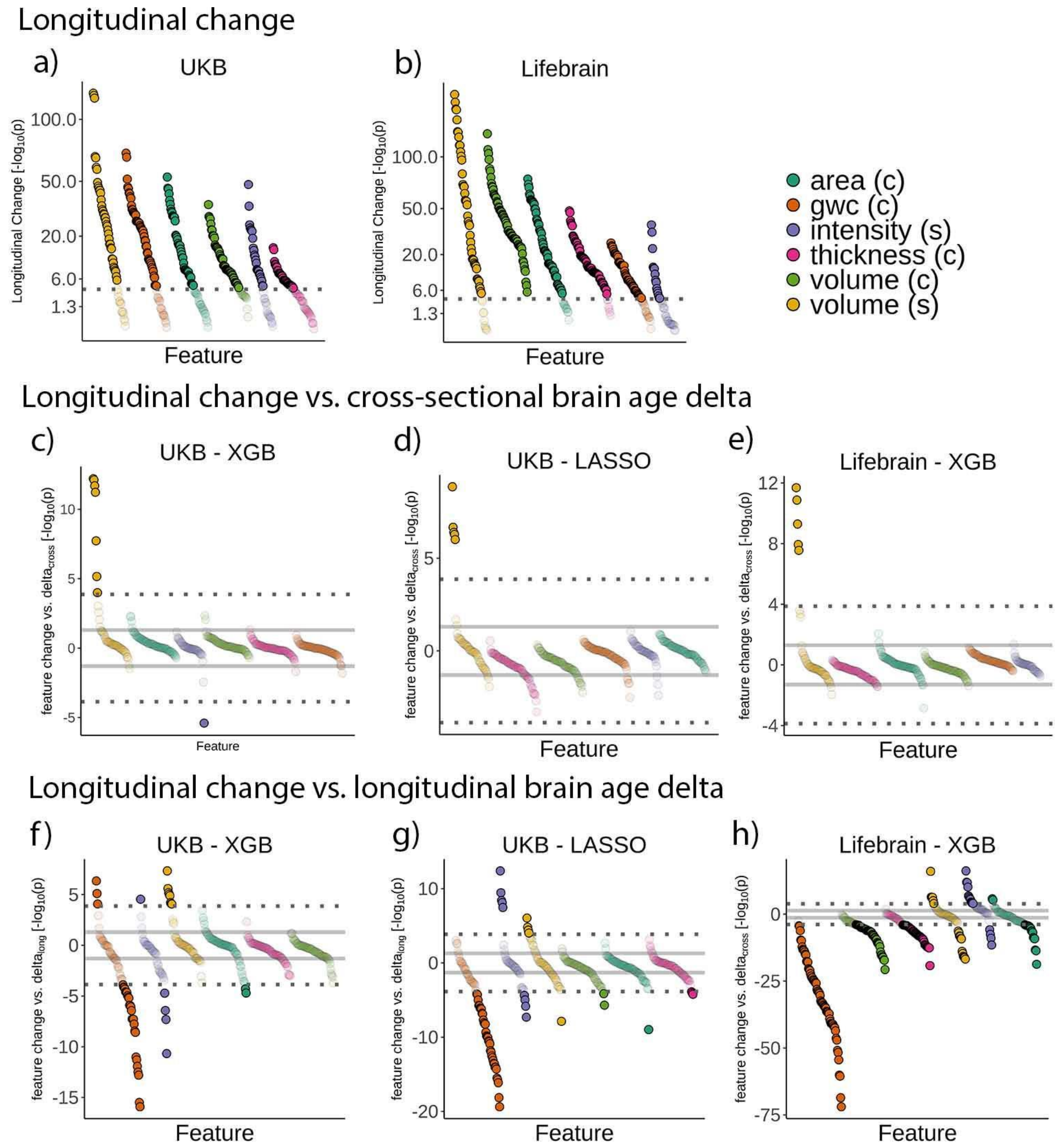

**Supplementary Fig. 4. Relationship between *brain age delta* and change in raw features.** Feature change over time in the **a)** UK Biobank and **b)** Lifebrain datasets. Signed relationship between cross-sectional *brain age delta* and longitudinal change in the raw features in **c)** the UK Biobank using a boosting gradient algorithm, **d)** the UK Biobank using a LASSO algorithm and **e)** the Lifebrain dataset using the boosting gradient algorithm. Signed relationship between change in *brain age delta (brain age* $delta_{long}$*)* and longitudinal change in the raw features in **f)** the UK Biobank using a boosting gradient algorithm, **g)** the UK Biobank using a LASSO algorithm and **h)** the Lifebrain dataset using the boosting gradient algorithm. Dashed lines represent a Bonferroni-corrected significance threshold (|n| = 365 and 372 features for UK Biobank and Lifebrain datasets, respectively. The solid line represents an uncorrected p = 0.05 significance threshold. n = 1,372 and 1,500 for the UK Biobank and the Lifebrain datasets.

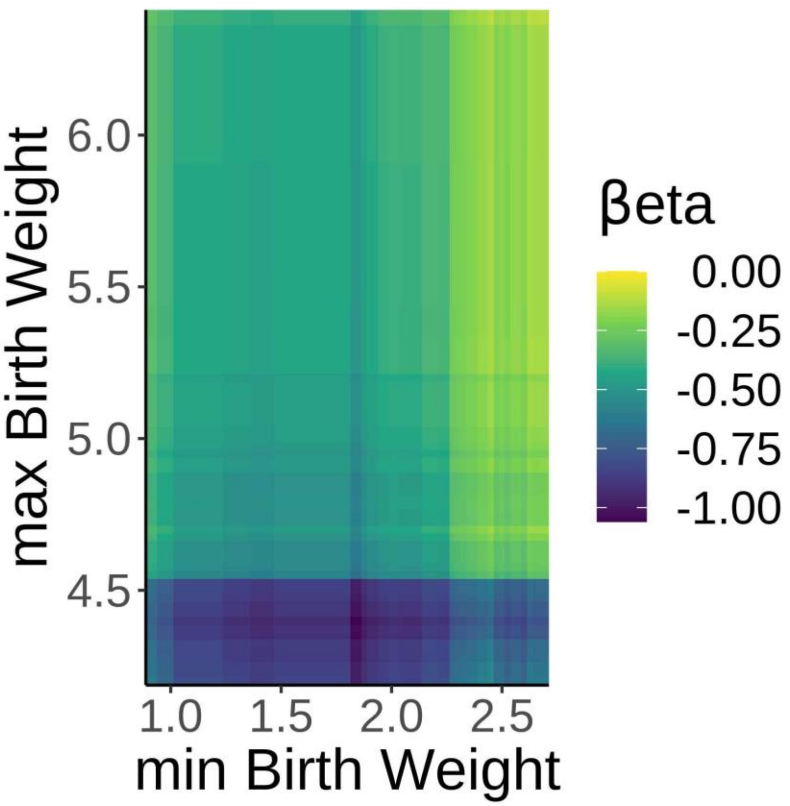

**Supplementary Fig. 5. Robust effects of birth weight on *brain age delta*.** βeta-estimates showing the relationship between *brain age delta* and birth weight with variable minimum and maximum birth weight exclusion thresholds. Note negative βetas irrespective of the minimum and maximum self-reported birth-weight thresholds.

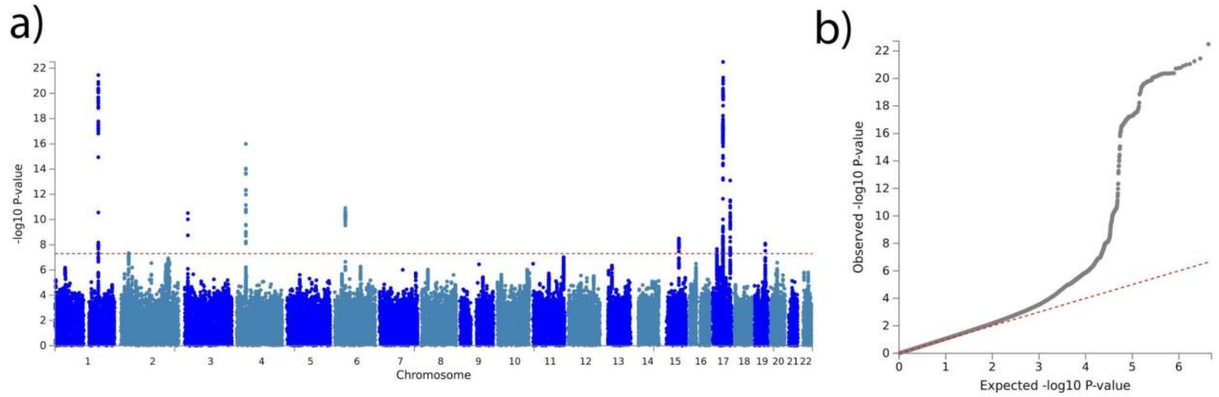

**Supplementary Fig. 6. Brain age delta GWAS. a)** Manhattan plot of the GWAS results for the test set on *brain age delta* (38,163 individuals). The horizontal line represents the threshold for genome-wide significance. **b)** Quantile-quantile (QQ) plot illustrating the deviation of the observed p-values from the null hypothesis.

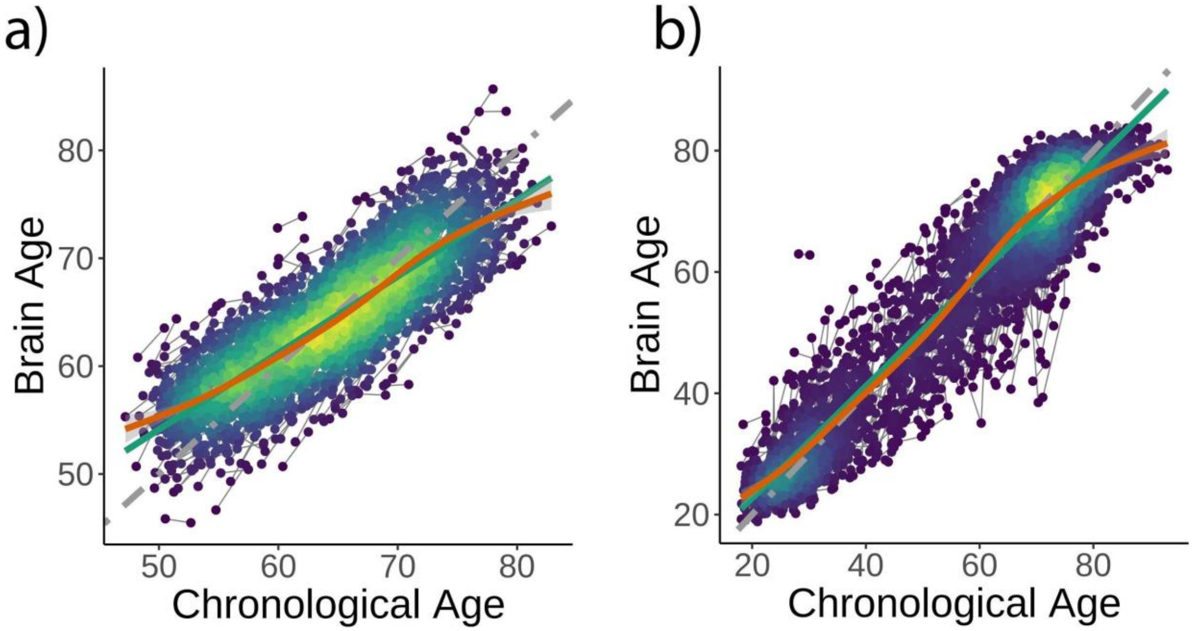

**Supplementary Fig. 7. Brain age model predictions.** *Brain age* model prediction (i.e. on test data) as estimated **a)** using LASSO in the UK Biobank dataset and **b)** extreme boosting gradient in the Lifebrain sample. Grey, green, and orange lines represent the identity, the linear, and the GAM functions fitting brain on chronological age.

| | GWC | Vol | Area | Cth | GWC | Vol | Area | Cth | GWC | Vol | Area | Cth | GWC | Vol | Area | Cth |
|---|---|---|---|---|---|---|---|---|---|---|---|---|---|---|---|---|
| | **UB Biobank** | | | | | | | | **Lifebrain** | | | | | | | |
| | **Left hemisphere** | | | | **Right hemisphere** | | | | **Left hemisphere** | | | | **Right hemisphere** | | | |
| **Total surface** | -- | -- | .00 | .10 | -- | .. | .00 | .10 | | | .17 | .60 | | | .16 | .58 |
| **Cingulate, caudal ant** | .06 | .01 | .00 | .01 | .07 | .01 | .00 | .00 | .38 | .09 | .02 | .09 | .40 | .13 | .05 | .11 |
| **Cingulate, rostral ant** | .20 | .01 | .00 | .02 | .15 | .01 | .00 | .00 | .52 | .16 | .03 | .26 | .44 | .14 | .06 | .12 |
| **Cingulate, posterior** | .13 | .01 | .00 | .02 | .16 | .02 | .01 | .03 | .48 | .23 | .09 | .30 | .52 | .22 | .10 | .27 |
| **Cingulate, isthmus** | .06 | .00 | .00 | .03 | .04 | .00 | .00 | .02 | .50 | .14 | .01 | .29 | .48 | .14 | .02 | .25 |
| **Insula** | .09 | .00 | .00 | .02 | .10 | .00 | .00 | .02 | .39 | .17 | .00 | .42 | .40 | .15 | .00 | .44 |
| **Frontal, superior** | .26 | .05 | .00 | .14 | .27 | .04 | .00 | .13 | .65 | .40 | .11 | .53 | .66 | .37 | .12 | .45 |
| **Frontal, caudal middle** | .18 | .03 | .00 | .08 | .20 | .02 | .00 | .07 | .60 | .23 | .06 | .42 | .61 | .20 | .06 | .39 |
| **Frontal, rostral middle** | .22 | .04 | .01 | .11 | .23 | .04 | .01 | .08 | .61 | .45 | .16 | .43 | .60 | .35 | .14 | .34 |
| **Frontal, pars opercularis** | .20 | .03 | .01 | .07 | .22 | .03 | .00 | .06 | .61 | .30 | .10 | .51 | .63 | .30 | .09 | .48 |
| **Frontal, pars triangularis** | .18 | .04 | .01 | .07 | .21 | .04 | .01 | .08 | .62 | .34 | .12 | .47 | .63 | .31 | .10 | .46 |
| **Frontal, pars orbitalis** | .16 | .04 | .01 | .04 | .19 | .05 | .01 | .04 | .56 | .37 | .16 | .26 | .58 | .38 | .16 | .26 |

| | | | | | | | | | | | | | | | | |
|---|---|---|---|---|---|---|---|---|---|---|---|---|---|---|---|---|
| **Frontal, lateral orbital** | .18 | .03 | .02 | .02 | .16 | .03 | .01 | .01 | .54 | .39 | .18 | .32 | .49 | .31 | .11 | .19 |
| **Frontal, medial orbital** | .20 | .02 | .00 | .02 | .22 | .03 | .00 | .01 | .53 | .19 | .03 | .27 | .53 | .24 | .08 | .16 |
| **Frontal, pole** | .19 | .02 | .01 | .03 | .17 | .01 | .01 | .02 | .53 | .27 | .14 | .12 | .47 | .21 | .09 | .07 |
| **Frontal, precentral gyrus** | .13 | .04 | .00 | .09 | .15 | .04 | .00 | .10 | .61 | .25 | .01 | .45 | .63 | .26 | .01 | .45 |
| **Parietal, postcentral gyrus** | .09 | .02 | .00 | .06 | .09 | .02 | .00 | .06 | .53 | .20 | .03 | .37 | .52 | .19 | .02 | .36 |
| **Parietal, paracentral gyrus** | .12 | .03 | .00 | .07 | .13 | .03 | .00 | .07 | .54 | .17 | .02 | .30 | .57 | .15 | .01 | .26 |
| **Parietal, superior** | .15 | .03 | .00 | .07 | .13 | .04 | .00 | .08 | .53 | .26 | .10 | .30 | .53 | .29 | .11 | .32 |
| **Parietal, inferior** | .16 | .03 | .00 | .08 | .14 | .04 | .00 | .11 | .53 | .29 | .12 | .40 | .54 | .32 | .11 | .46 |
| **Parietal, supramarginal** | .16 | .02 | .00 | .09 | .18 | .02 | .00 | .09 | .55 | .27 | .08 | .53 | .57 | .28 | .05 | .54 |
| **Parietal, precuneus** | .20 | .05 | .00 | .09 | .18 | .03 | .00 | .08 | .57 | .31 | .11 | .47 | .56 | .28 | .10 | .43 |
| **Temporal, parahippocampal** | .06 | .03 | .02 | .02 | .06 | .02 | .01 | .02 | .24 | .15 | .12 | .06 | .32 | .16 | 8.0 | .10 |
| **Temporal, entorhinal** | .09 | .00 | .00 | .04 | .09 | .00 | .00 | .03 | .25 | .02 | .01 | .07 | .25 | .01 | .00 | .07 |
| **Temporal, pole** | .10 | -- | -- | -- | .11 | -- | -- | -- | .31 | .06 | .05 | .07 | .29 | .12 | .03 | .10 |
| **Temporal, superior** | .15 | .04 | .00 | .10 | .19 | .05 | .00 | .11 | .53 | .34 | .09 | .56 | .57 | .36 | .11 | .56 |
| **Temporal, middle** | .17 | .04 | .01 | .04 | .19 | .04 | .01 | .05 | .51 | .40 | .19 | .49 | .54 | .40 | .19 | .48 |

| | | | | | | | | | | | | | | | | |
|---|---|---|---|---|---|---|---|---|---|---|---|---|---|---|---|---|
| **Temporal, inferior** | .17 | .02 | .02 | .02 | .17 | .02 | .01 | .03 | .48 | .28 | .18 | .35 | .48 | .27 | .16 | .32 |
| **Temporal, transverse** | .02 | .00 | .01 | .00 | .03 | .00 | .01 | .00 | .47 | .17 | .06 | .27 | .47 | .20 | .17 | .29 |
| **Temporal, bank sup temp sulc** | .09 | .02 | .01 | .03 | .13 | .02 | .01 | .03 | .47 | .19 | .08 | .32 | .49 | .28 | .13 | .36 |
| **Temporal, fusiform** | .17 | .03 | .01 | .05 | .16 | .03 | .01 | .06 | .48 | .27 | .19 | .30 | .48 | .23 | .14 | .31 |
| **Occipital, lateral** | .03 | .02 | .01 | .03 | .02 | .03 | .01 | .02 | .36 | .25 | .15 | .21 | .35 | .21 | .11 | .25 |
| **Occipital, pericalcarine** | .00 | .00 | .00 | .00 | .02 | .00 | .00 | .00 | .27 | .06 | .03 | .18 | .22 | .06 | .04 | .13 |
| **Occipital, lingual** | .00 | .01 | .01 | .01 | .00 | .01 | .01 | .00 | .27 | .18 | .09 | .30 | .26 | .19 | .09 | .30 |
| **Occipital, cuneus** | .00 | .01 | .01 | .00 | .00 | .00 | .01 | .00 | .27 | .12 | .08 | .18 | .27 | .12 | .08 | .19 |

**Supplementary Table 1. List of cortical brain features.** List of cortical features included in the brain age model and age variance explained in the UK Biobank and the Lifebrain training datasets. Vol = volume. GWC = Gray-white matter contrast. Cth = Cortical Thickness.

| | Vol | Int | Vol | Int | Vol | Int | Vol | Int | Vol | Int | Vol | Int |
|---|---|---|---|---|---|---|---|---|---|---|---|---|
| | UK Biobank | | | | | | Lifebrain | | | | | |
| | Left hemi | | Right hemi | | Bilateral | | Left hemi | | Right hemi | | Bilateral | |
| **Ventricular ROIs** | | | | | | | | | | | | |
| **3rd Ventricle** | -- | -- | -- | -- | .19 | .22 | -- | -- | -- | -- | .55 | .45 |
| **4th Ventricle** | -- | -- | -- | -- | .02 | .02 | -- | -- | -- | -- | .02 | .01 |
| **5th Ventricle** | -- | -- | -- | -- | .00 | .00 | -- | -- | -- | -- | .05 | .03 |
| **Inf lat vent** | .18 | .12 | .15 | .06 | -- | -- | .41 | .11 | .39 | .02 | -- | -- |
| **Lat vent** | .16 | .11 | .16 | .11 | -- | -- | .43 | .11 | .45 | .12 | -- | -- |
| **CSF total** | -- | -- | -- | -- | .06 | .12 | -- | -- | -- | -- | .21 | .23 |
| **Volumetric ROIs** | | | | | | | | | | | | |
| **Accumbens** | .18 | .11 | .12 | .15 | -- | -- | .45 | .31 | .41 | .38 | -- | -- |
| **Amygdala** | .11 | .01 | .06 | .05 | -- | -- | .33 | .21 | .25 | .31 | -- | -- |
| **Brainstem** | -- | -- | -- | -- | .00 | .13 | -- | -- | -- | -- | .07 | .06 |
| **Caudate** | .00 | .01 | .00 | .01 | -- | -- | .18 | .22 | .16 | .15 | -- | -- |
| **Cerebellum** | .01 | .01 | .01 | .01 | -- | -- | .35 | .02 | .32 | .01 | -- | -- |

| | | | | | | | | | | | | |
|---|---|---|---|---|---|---|---|---|---|---|---|---|
| **Hippocampus** | .11 | .02 | .11 | .02 | -- | -- | .37 | .24 | .39 | .25 | -- | -- |
| **Pallidum** | .01 | .04 | .01 | .03 | -- | -- | .09 | .10 | .04 | .12 | -- | -- |
| **Putamen** | .04 | .01 | .04 | .00 | -- | -- | .35 | .13 | .37 | .12 | -- | -- |
| **Thalamus** | .10 | .15 | .07 | .16 | -- | -- | .45 | .04 | .43 | .04 | -- | -- |
| **White matter** | | | | | | | | | | | | |
| **CC anterior** | -- | -- | -- | -- | .02 | .06 | -- | -- | -- | -- | .10 | .10 |
| **CC central** | -- | -- | -- | -- | .08 | .05 | -- | -- | -- | -- | .28 | .04 |
| **CC mid anterior** | -- | -- | -- | -- | .10 | .04 | -- | -- | -- | -- | .22 | .06 |
| **CC mid posterior** | -- | -- | -- | -- | .05 | .13 | -- | -- | -- | -- | .24 | .12 |
| **CC posterior** | -- | -- | -- | -- | .00 | .07 | -- | -- | -- | -- | .03 | .08 |
| **Cerebellum WM** | .07 | .00 | .06 | .00 | -- | -- | .08 | .02 | .10 | .02 | -- | -- |
| **Cerebral WM** | .03 | -- | .03 | -- | -- | -- | .15 | -- | .13 | -- | .14 | -- |
| **WM hypointensities** | -- | -- | -- | -- | .14 | .05 | -- | -- | -- | -- | .31 | .15 |
| **Non WM hypointensities** | -- | -- | -- | -- | .00 | .00 | -- | -- | -- | -- | .21 | .00 |
| **Global features** | | | | | | | | | | | | |
| **Brain Seg** | -- | -- | -- | -- | .02 | -- | -- | -- | -- | -- | .23 | -- |
| **Brain Seg not vent** | -- | -- | -- | -- | .05 | -- | -- | -- | -- | -- | .32 | -- |

| | | | | | | | | | | | | |
|---|---|---|---|---|---|---|---|---|---|---|---|---|
| **Brain Seg not vent surf** | -- | -- | -- | -- | .05 | -- | -- | -- | -- | -- | .32 | -- |
| **Cortex** | .05 | -- | .05 | -- | -- | -- | .45 | -- | .44 | -- | .45 | -- |
| **eICV** | -- | -- | -- | -- | .00 | -- | -- | -- | -- | -- | .00 | -- |
| **Subcort gray** | -- | -- | -- | -- | .06 | -- | -- | -- | -- | -- | .44 | -- |
| **Supratentorial** | -- | -- | -- | -- | .02 | -- | -- | -- | -- | -- | .21 | -- |
| **Supratentorial not vent** | -- | -- | -- | -- | .05 | -- | -- | -- | -- | -- | .30 | -- |
| **Total gray** | -- | -- | -- | -- | .05 | -- | -- | -- | -- | -- | .47 | -- |
| **Other** | | | | | | | | | | | | |
| **Choroid plexus** | .18 | .17 | .18 | .18 | -- | -- | .35 | .33 | .34 | .32 | -- | -- |
| **Optic chiasm** | -- | -- | -- | -- | .05 | .00 | -- | -- | -- | -- | .09 | .01 |
| **Ventral DC** | .05 | .06 | .06 | .05 | -- | -- | .19 | .11 | .25 | .09 | -- | -- |
| **Ventricle choroid** | -- | -- | -- | -- | .18 | -- | -- | -- | -- | -- | -- | -- |
| **Vessel** | .00 | .00 | .01 | .00 | -- | -- | .02 | .02 | .03 | .01 | -- | -- |

**Supplementary Table 2. List of subcortical brain features.** List of subcortical features included in the brain age model and age variance explained in the UK Biobank and the Lifebrain training datasets. Vol = volume; Int = Intensity; hemi = hemisphere.

| | Training dataset | | | | Test dataset | | | | |
|---|---|---|---|---|---|---|---|---|---|
| **Cohort** | N | Age | Age Range | Sex (f:m) | N (Obs) | Age | Age Range | Sex (f:m) | Follow-up |
| **UK Biobank** | 38,682 | 64.4 (7.6) | 44.8 - 82.6 | 20,470:18,212 | 1,372 (2) | 63.4 (7.2) | 47.2 - 80.6 | 685:687 | 2.3 (.1) |
| **Lifebrain Total** | 1792 | 50.3 (21.8) | 18.0 - 94.4 | 1,075:717 | 1,500 (2.8 [1.2]) | 56.9 (18.9) | 18.1 - 89.0 | 769:731 | 3.4 (2.2) |
| LCBC | 838 | 35.5 (16.5) | 18.0 - 93.4 | 563:275 | 556 (3.7 [1.5]) | 48.9 (20.7) | 18.1 - 85.4 | 1,229:848 | 4.8 (2.7) |
| Cam-CAN | 386 | 54.7 (18.7) | 18.6 - 87.4 | 196:190 | 255 (2) | 55.0 (18.1) | 19.3 - 89.0 | 262:248 | 1.4 (0.7) |
| Base-II | 126 | 59.8 (18.1) | 25.0 - 82.0 | 58:68 | 319 (2) | 62.0 (16.6) | 24.1 - 81.3 | 224:414 | 1.9 (0.7) |
| Betula | 139 | 64.8 (13.0) | 25.9 - 84.6 | 75:64 | 170 (2) | 59.8 (13.8) | 25.5 - 80.8 | 166:174 | 4.0 (0.2) |
| UB | 6 | 64.3 (11.8) | 43.5 - 77.7 | 2:4 | 80 (2.7 [.4]) | 67.3 (6.9) | 36.8 - 78.1 | 135:84 | 3.7 (0.9) |
| AIBL* | 297 | 75.1 (5.7) | 61.8 - 94.4 | 181:116 | 120 (3.4 [.8]) | 73.0 (7.0) | 62.6 - 88.4 | 214:195 | 4.0 (1.4) |

**Supplementary Table 3. Sociodemographics.** Main sample descriptives for the training and test datasets. Obs = Mean number of observations per participant [SD]. Follow-up = Mean time (years) between the first and the last MRI observation (SD). For the test datasets, Age and Age Range refers to Age at baseline. *AIBL does not belong to the Lifebrain consortium but was included to enrich the replication sample.

| | | | UK Biobank | | | | | | Lifebrain | | | |
|---|---|---|---|---|---|---|---|---|---|---|---|---|
| **Modality** | **Region** | **Hemi** | **Long. change** | **$PC_1$ load** | ***$delta_{lross}$* - XGB** | ***$delta_{cross}$* - LASSO** | **$delta_{long}$ - XGB** | **$delta_{long}$ - LASSO** | **Long. change** | **$PC_1$ load** | ***$delta_{cross}$*** | ***$delta_{long}$*** |
| **Area** | **Total surface** | Left | -.03 (45.5) | 0.012 | .001 (0.8) | .000 (0.1) | .001 (-0.5) | .002 (-1.1) | -.03 (68.3) | 0.005 | .000 (0.4) | .007 (3.0) |
| | | Right | -.03 (52.9) | 0.004 | .002 (0.8) | .000 (0.2) | .000 (-0.1) | .006 (-2.5) | -.03 (57.0) | 0.006 | .000 (-0.2) | .007 (-2.9) |
| | **Cingulate, caudal ant** | Left | -.01 (1.5) | -- | .003 (1.2) | .000 (0.2) | .000 (0.3) | .000 (0.0) | -.01 (14.6) | 0.009 | .000 (-0.1) | .025 (-9.1) |
| | | Right | -.01 (1.0) | -- | .000 (0.0) | .001 (0.8) | .008 (-3.2) | .001 (-0.7) | -.02 (31.9) | 0.008 | .002 (-1.1) | .010 (-3.9) |
| | **Cingulate, rostral ant** | Left | -.01 (1.2) | -- | .000 (-0.3) | .000 (-0.2) | .000 (0.0) | .000 (0.3) | -.02 (10.3) | 0.008 | .001 (0.9) | .009 (3.7) |
| | | Right | -.01 (3.0) | -- | .001 (0.7) | .002 (0.9) | .010 (-3.7) | .001 (-0.6) | -.01 (5.9) | 0.011 | .002 (-0.9) | .011 (-4.3) |
| | **Cingulate, posterior** | Left | -.02 (4.8) | 0.016 | .001 (0.7) | .000 (0.0) | .003 (1.4) | .001 (0.5) | -.03 (49.8) | 0.011 | .000 (0.0) | .011 (4.2) |
| | | Right | -.01 (3.4) | -- | .000 (0.1) | .001 (0.5) | .007 (-2.9) | .001 (-0.6) | -.03 (76.5) | 0.011 | .002 (-0.9) | .010 (-4.1) |
| | **Cingulate, isthmus** | Left | -.01 (2.0) | -- | .000 (0.0) | .000 (0.2) | .000 (0.2) | .005 (1.9) | -.02 (10.9) | 0.003 | .001 (-0.6) | .001 (-0.7) |
| | | Right | -.01 (2.5) | -- | .000 (0.4) | .000 (0.3) | .001 (0.8) | .001 (0.5) | -.02 (10.3) | 0.011 | .000 (0.2) | .005 (2.3) |
| | **Insula** | Left | -.00 (0.0) | -- | .000 (0.3) | .000 (0.0) | .002 (-0.9) | .001 (-0.8) | -.01 (2.9) | -- | .000 (-0.2) | .004 (-1.7) |
| | | Right | -.02 (1.8) | -- | .001 (-0.5) | .000 (-0.2) | .000 (-0.1) | .000 (-0.1) | -.00 (0.6) | -- | .000 (-0.3) | .005 (-2.1) |
| | **Frontal, superior** | Left | -.01 (7.9) | 0.006 | .005 (1.9) | .001 (0.5) | .000 (-0.2) | .000 (-0.3) | -.02 (15.3) | -0.001 | .000 (0.2) | .008 (3.2) |
| | | Right | -.01 (3.8) | -- | .006 (2.3) | .001 (0.5) | .002 (-0.9) | .001 (-0.4) | -.01 (14.7) | -0.001 | .000 (0.0) | .011 (4.5) |
| | **Frontal, caudal middle** | Left | -.01 (3.2) | -- | .002 (0.9) | .001 (0.4) | .001 (0.6) | .000 (0.1) | -.01 (7.7) | -0.006 | .000 (-0.1) | .001 (-0.9) |
| | | Right | -.01 (1.5) | -- | .001 (0.6) | .000 (0.2) | .000 (-0.2) | .002 (-1.0) | -.01 (6.8) | -0.004 | .000 (-0.4) | .001 (-0.6) |
| | **Frontal, rostral middle** | Left | -.02 (11.8) | 0.011 | .001 (0.6) | .000 (0.3) | .000 (0.0) | .003 (1.4) | -.02 (22.1) | 0.006 | .000 (-0.1) | .009 (-3.6) |
| | | Right | -.01 (3.2) | -- | .002 (1.1) | .000 (0.1) | .005 (-2.1) | .002 (-1.2) | -.01 (12.8) | 0.009 | .001 (-0.7) | .037 (-13.4) |

| | | | | | | | | | | | | |
|---|---|---|---|---|---|---|---|---|---|---|---|---|
| | **Frontal, pars opercularis** | Left | -.02 (8.3) | 0.007 | .001 (0.7) | .000 (0.3) | .000 (0.4) | .000 (0.4) | -.02 (26.5) | 0.001 | .000 (-0.3) | .001 (-0.6) |
| | | Right | -.02 (6.6) | 0.002 | .000 (0.1) | .001 (0.7) | .001 (0.5) | .002 (0.9) | -.01 (12.5) | 0.002 | .000 (0.0) | .006 (2.6) |
| | **Frontal, pars triangularis** | Left | -.04 (16.4) | 0.010 | .000 (0.3) | .000 (0.0) | .000 (0.0) | .002 (1.0) | -.02 (14.7) | 0.005 | .000 (0.4) | .009 (3.6) |
| | | Right | -.02 (8.2) | 0.003 | .000 (0.3) | .001 (0.5) | .005 (-2.0) | .007 (-2.6) | -.02 (10.3) | 0.011 | .000 (-0.1) | .017 (-6.2) |
| | **Frontal, pars orbitalis** | Left | -.03 (18.5) | 0.021 | .000 (0.1) | .000 (0.1) | .002 (1.0) | .002 (0.9) | -.02 (34.4) | 0.001 | .001 (0.4) | .002 (0.9) |
| | | Right | -.02 (6.6) | 0.004 | .001 (0.5) | .000 (0.1) | .000 (0.1) | .000 (0.2) | -.02 (23.7) | 0.004 | .000 (-0.2) | .005 (-2.3) |
| | **Frontal, lateral orbital** | Left | -.03 (10.2) | 0.016 | .000 (0.0) | .000 (0.0) | .013 (-4.7) | .026 (-9.0) | -.02 (15.4) | 0.019 | .001 (-0.7) | .024 (-8.7) |
| | | Right | -.02 (1.5) | -- | .001 (-0.4) | .000 (-0.3) | .009 (-3.5) | .009 (-3.5) | -.01 (2.0) | -- | .000 (-0.1) | .003 (-1.3) |
| | **Frontal, medial orbital** | Left | -.02 (2.9) | -- | .000 (0.0) | .001 (0.4) | .000 (0.1) | .001 (0.4) | -.02 (2.7) | -- | .000 (0.1) | .006 (2.6) |
| | | Right | -.01 (1.1) | -- | .000 (0.0) | .000 (0.0) | .001 (-0.7) | .003 (-1.3) | -.01 (1.5) | -- | .000 (-0.2) | .014 (-5.4) |
| | **Frontal, pole** | Left | -.04 (7.7) | 0.019 | .000 (0.1) | .001 (0.6) | .000 (0.0) | .002 (0.9) | -.06 (15.7) | 0.027 | .000 (-0.2) | .018 (-6.9) |
| | | Right | -.01 (0.7) | -- | .000 (-0.2) | .001 (-0.7) | .001 (0.7) | .001 (0.5) | -.04 (7.9) | 0.021 | .000 (0.0) | .007 (3.1) |
| | **Frontal, precentral gyrus** | Left | -.01 (3.2) | -- | .001 (0.5) | .000 (0.2) | .000 (0.0) | .002 (0.9) | -.01 (1.5) | -- | .001 (0.4) | .005 (2.2) |
| | | Right | -.00 (0.7) | -- | .001 (0.5) | .000 (0.3) | .000 (0.1) | .007 (2.7) | -.01 (2.0) | -- | .000 (0.1) | .015 (5.7) |
| | **Parietal, postcentral gyrus** | Left | -.01 (8.4) | -0.004 | .000 (0.2) | .000 (0.1) | .000 (0.1) | .001 (0.6) | -.01 (6.7) | -0.014 | .001 (0.5) | .005 (2.1) |
| | | Right | -.01 (5.0) | -0.007 | .001 (-0.7) | .000 (-0.2) | .005 (2.1) | .001 (0.7) | -.01 (3.4) | -- | .001 (0.5) | .014 (5.3) |
| | **Parietal, paracentral gyrus** | Left | -.00 (0.3) | -- | .002 (0.9) | .002 (0.9) | .001 (0.6) | .001 (0.8) | -.01 (2.8) | -- | .001 (0.6) | .001 (0.8) |
| | | Right | -.01 (1.0) | -- | .000 (-0.1) | .000 (-0.4) | .001 (-0.7) | .000 (-0.3) | -.01 (2.1) | -- | .000 (0.0) | .007 (3.0) |
| | **Parietal, superior** | Left | -.02 (20.4) | 0.005 | .000 (-0.2) | .002 (-1.1) | .000 (0.2) | .005 (1.9) | -.02 (21.8) | -0.002 | .000 (-0.2) | .001 (-0.6) |
| | | Right | -.02 (8.0) | 0.005 | .001 (0.6) | .000 (0.1) | .007 (2.7) | .000 (0.2) | -.02 (20.4) | -0.004 | .000 (-0.1) | .001 (-0.6) |
| | **Parietal, inferior** | Left | -.03 (33.1) | 0.017 | .001 (0.5) | .002 (0.9) | .001 (0.7) | .004 (1.8) | -.02 (62.1) | 0.005 | .000 (-0.1) | .006 (-2.6) |
| | | Right | -.03 (29.0) | 0.004 | .000 (0.2) | .000 (0.3) | .000 (-0.2) | .000 (-0.2) | -.02 (53.3) | 0.003 | .000 (0.0) | .000 (0.1) |
| | **Parietal, supramarginal** | Left | -.02 (13.9) | 0.008 | .000 (-0.1) | .001 (-0.8) | .001 (0.4) | .001 (0.6) | -.02 (40.8) | 0.001 | .000 (0.2) | .002 (1.2) |
| | | Right | -.01 (4.6) | -0.005 | .001 (-0.6) | .000 (-0.1) | .000 (0.3) | .000 (0.2) | -.02 (33.1) | -0.002 | .000 (0.0) | .001 (0.6) |

| | | | | | | | | | | | | |
|---|---|---|---|---|---|---|---|---|---|---|---|---|
| | **Parietal, precuneus** | Left | -.02 (17.1) | 0.013 | .001 (0.4) | .001 (0.6) | .006 (2.5) | .001 (0.6) | -.02 (55.8) | 0.004 | .000 (0.4) | .004 (1.7) |
| | | Right | -.02 (12.9) | 0.013 | .002 (1.2) | .000 (0.2) | .009 (3.4) | .001 (0.6) | -.02 (49.4) | 0.001 | .000 (-0.2) | .000 (-0.3) |
| | **Temporal, parahippocampal** | Left | -.03 (6.9) | -0.009 | .001 (-0.5) | .001 (-0.4) | .004 (-1.6) | .004 (-1.8) | -.02 (14.1) | 0.004 | .005 (2.1) | .002 (0.9) |
| | | Right | -.03 (7.8) | 0.001 | .000 (0.0) | .000 (0.2) | .000 (0.0) | .000 (0.4) | -.02 (15.4) | 0.013 | .003 (-1.3) | .012 (-4.7) |
| | **Temporal, entorhinal** | Left | -.00 (0.2) | -- | .000 (0.2) | .001 (0.5) | .001 (0.5) | .000 (0.2) | -.02 (5.2) | 0.003 | .000 (0.3) | .001 (0.4) |
| | | Right | -.01 (0.4) | -- | .002 (1.1) | .002 (0.9) | .001 (0.4) | .000 (0.0) | -.01 (2.5) | -- | .001 (-0.5) | .008 (-3.5) |
| | **Temporal, pole** | Left | -- | -- | -- | -- | -- | -- | -.03 (6.0) | 0.021 | .001 (0.5) | .001 (0.6) |
| | | Right | -- | -- | -- | -- | -- | -- | -.03 (7.3) | 0.035 | .000 (-0.1) | .007 (-2.9) |
| | **Temporal, superior** | Left | -.02 (16.2) | 0.009 | .000 (0.3) | .000 (0.2) | .000 (-0.1) | .003 (-1.2) | -.02 (56.6) | 0.006 | .000 (0.2) | .000 (0.2) |
| | | Right | -.02 (20.0) | -0.005 | .000 (-0.1) | .000 (-0.2) | .000 (-0.1) | .002 (-0.9) | -.02 (33.4) | 0.004 | .001 (-0.8) | .000 (-0.1) |
| | **Temporal, middle** | Left | -.04 (45.0) | 0.012 | .000 (0.1) | .000 (0.1) | .002 (-1.0) | .001 (-0.5) | -.03 (71.2) | 0.012 | .001 (0.5) | .011 (4.3) |
| | | Right | -.03 (38.0) | 0.007 | .000 (-0.3) | .000 (-0.3) | .006 (-2.6) | .003 (-1.3) | -.03 (60.7) | 0.011 | .000 (-0.1) | .006 (-2.8) |
| | **Temporal, inferior** | Left | -.04 (42.0) | 0.015 | .000 (-0.1) | .000 (-0.4) | .012 (-4.3) | .009 (-3.5) | -.02 (46.5) | 0.017 | .000 (-0.3) | .052 (-18.7) |
| | | Right | -.03 (31.2) | 0.007 | .000 (0.1) | .001 (0.6) | .003 (-1.3) | .007 (-2.7) | -.03 (54.4) | 0.015 | .000 (-0.2) | .038 (-13.7) |
| | **Temporal, transverse** | Left | -.02 (4.8) | 0.005 | .001 (0.4) | .001 (0.4) | .004 (1.6) | .000 (0.0) | -.03 (18.4) | 0.007 | .000 (0.3) | .001 (0.6) |
| | | Right | -.01 (0.8) | -- | .000 (-0.4) | .001 (-0.5) | .000 (0.1) | .000 (0.3) | -.05 (25.5) | 0.001 | .000 (-0.1) | .001 (-0.5) |
| | **Temporal, bank sup temp sulc** | Left | -.03 (19.4) | 0.007 | .000 (0.0) | .000 (0.0) | .000 (-0.2) | .001 (-0.7) | -.01 (21.5) | 0.001 | .002 (-1.3) | .005 (-2.3) |
| | | Right | -.03 (16.4) | -0.003 | .000 (0.0) | .000 (0.2) | .001 (-0.6) | .000 (-0.3) | -.02 (37.9) | 0.002 | .006 (-2.9) | .001 (-0.8) |
| | **Temporal, fusiform** | Left | -.04 (31.8) | 0.013 | .001 (0.7) | .001 (0.7) | .003 (-1.5) | .006 (-2.4) | -.03 (55.2) | 0.014 | .000 (0.0) | .005 (2.4) |
| | | Right | -.04 (28.8) | 0.017 | .003 (1.5) | .001 (0.5) | .000 (-0.3) | .006 (-2.4) | -.02 (38.6) | 0.014 | .000 (-0.1) | .011 (-4.4) |
| | **Occipital, lateral** | Left | -.03 (29.5) | 0.021 | .000 (0.3) | .000 (0.2) | .003 (1.3) | .007 (2.7) | -.03 (57.4) | 0.009 | .000 (0.4) | .000 (0.3) |

| | | | | | | | | | | | | |
|---|---|---|---|---|---|---|---|---|---|---|---|---|
| | | Right | -.03 (35.7) | 0.015 | .000 (0.3) | .001 (0.6) | .002 (1.1) | .003 (1.5) | -.03 (41.8) | 0.009 | .000 (0.3) | .002 (0.9) |
| | **Occipital, pericalcarine** | Left | -.01 (6.4) | 0.017 | .006 (2.3) | .001 (0.4) | .000 (0.3) | .003 (1.4) | -.02 (20.1) | 0.008 | .002 (1.0) | .001 (0.7) |
| | | Right | -.02 (11.9) | 0.013 | .002 (0.8) | .000 (0.2) | .001 (0.5) | .000 (0.2) | -.01 (6.7) | 0.008 | .003 (1.5) | .003 (1.4) |
| | **Occipital, lingual** | Left | -.03 (29.3) | 0.016 | .000 (0.3) | .000 (0.2) | .010 (-3.6) | .006 (-2.4) | -.03 (38.9) | 0.012 | .002 (1.1) | .017 (6.4) |
| | | Right | -.03 (27.0) | 0.018 | .000 (-0.2) | .002 (-1.0) | .001 (0.5) | .001 (0.7) | -.03 (28.6) | 0.014 | .000 (0.0) | .025 (9.2) |
| | **Occipital, cuneus** | Left | -.02 (14.9) | 0.016 | .000 (0.1) | .001 (0.6) | .003 (1.2) | .002 (0.8) | -.02 (20.7) | 0.009 | .001 (0.8) | .013 (5.0) |
| | | Right | -.02 (8.8) | 0.017 | .000 (-0.2) | .002 (-1.0) | .002 (1.0) | .003 (1.4) | -.02 (12.5) | 0.010 | .000 (-0.3) | .003 (-1.3) |
| Cth | **Total surface** | Left | -.05 (8.2) | 0.149 | .000 (-0.1) | .001 (-0.8) | .004 (-1.8) | .005 (-2.0) | -.06 (40.7) | -0.001 | .002 (-1.0) | .033 (-12.2) |
| | | Right | -.04 (5.3) | 0.126 | .000 (0.0) | .004 (1.9) | .000 (-0.2) | .000 (-0.3) | -.07 (38.0) | 0.000 | .002 (-1.0) | .031 (-11.4) |
| | **Cingulate, caudal ant** | Left | -.01 (0.6) | -- | .001 (0.7) | .001 (0.5) | .000 (-0.3) | .002 (-1.0) | -.03 (9.4) | -0.026 | .000 (-0.4) | .000 (-0.0) |
| | | Right | -.02 (1.0) | -- | .000 (-0.2) | .002 (-1.2) | .000 (0.3) | .000 (0.3) | -.01 (1.1) | -- | .000 (-0.2) | .001 (-0.9) |
| | **Cingulate, rostral ant** | Left | -.04 (5.1) | 0.086 | .000 (0.0) | .000 (0.2) | .001 (-0.5) | .003 (-1.3) | -.03 (6.8) | -0.003 | .002 (-1.0) | .004 (-1.8) |
| | | Right | -.02 (1.0) | -- | .004 (-1.7) | .006 (-2.3) | .000 (0.2) | .000 (0.3) | -.01 (1.5) | -- | .000 (-0.2) | .004 (-1.8) |
| | **Cingulate, posterior** | Left | -.04 (4.1) | 0.088 | .001 (-0.7) | .002 (-1.0) | .000 (0.0) | .001 (0.6) | -.05 (27.2) | -0.015 | .002 (-1.0) | .001 (-0.5) |
| | | Right | -.02 (1.1) | -- | .000 (-0.2) | .002 (-0.9) | .000 (-0.1) | .000 (-0.4) | -.03 (11.6) | -0.019 | .000 (0.2) | .000 (0.3) |
| | **Cingulate, isthmus** | Left | -.02 (2.0) | -- | .000 (0.1) | .000 (0.1) | .002 (-1.0) | .002 (-1.1) | -.04 (26.4) | 0.007 | .000 (-0.1) | .022 (-8.2) |
| | | Right | -.01 (1.3) | -- | .001 (-0.4) | .005 (-2.0) | .000 (0.0) | .006 (2.3) | -.03 (13.7) | 0.005 | .000 (-0.1) | .024 (-8.6) |
| | **Insula** | Left | -.04 (3.7) | -- | .001 (-0.7) | .001 (-0.6) | .000 (-0.3) | .000 (-0.1) | -.05 (23.7) | 0.004 | .001 (-0.4) | .018 (-6.8) |
| | | Right | -.01 (0.5) | -- | .000 (0.0) | .000 (0.4) | .000 (-0.1) | .000 (-0.0) | -.05 (23.6) | -0.007 | .001 (-0.7) | .000 (-0.1) |
| | **Frontal, superior** | Left | -.05 (9.0) | 0.109 | .001 (-0.8) | .003 (-1.5) | .002 (-1.0) | .010 (-3.8) | -.05 (19.5) | -0.037 | .001 (-0.6) | .005 (-2.1) |

| | | | | | | | | | | | | |
|---|---|---|---|---|---|---|---|---|---|---|---|---|
| | | Right | -.04 (5.6) | 0.096 | .001 (-0.6) | .009 (-3.3) | .001 (0.5) | .003 (1.4) | -.04 (13.8) | -0.040 | .002 (-1.3) | .004 (-1.7) |
| | **Frontal, caudal middle** | Left | -.02 (1.8) | -- | .000 (0.0) | .001 (0.7) | .002 (-0.9) | .004 (-1.8) | -.06 (18.0) | -0.014 | .002 (-1.2) | .017 (-6.3) |
| | | Right | -.03 (3.0) | -- | .000 (-0.1) | .003 (-1.5) | .000 (0.1) | .000 (0.3) | -.05 (11.4) | -0.022 | .001 (-0.7) | .007 (-3.0) |
| | **Frontal, rostral middle** | Left | -.05 (10.7) | 0.109 | .001 (-0.5) | .003 (-1.4) | .003 (-1.2) | .011 (-4.2) | -.05 (12.7) | -0.023 | .002 (-1.0) | .015 (-5.8) |
| | | Right | -.04 (7.3) | 0.102 | .000 (-0.1) | .006 (-2.4) | .000 (-0.1) | .003 (-1.2) | -.05 (11.8) | -0.033 | .000 (-0.2) | .010 (-4.1) |
| | **Frontal, pars opercularis** | Left | -.04 (5.0) | 0.107 | .000 (0.0) | .002 (0.9) | .005 (-2.1) | .011 (-4.1) | -.04 (14.4) | -0.010 | .001 (-0.6) | .010 (-4.2) |
| | | Right | -.02 (3.0) | -- | .000 (0.2) | .001 (0.7) | .000 (-0.3) | .000 (-0.4) | -.05 (18.3) | -0.012 | .000 (-0.3) | .011 (-4.3) |
| | **Frontal, pars triangularis** | Left | -.02 (2.0) | -- | .000 (-0.2) | .001 (-0.6) | .002 (-0.9) | .005 (-2.2) | -.04 (9.6) | -0.010 | .000 (-0.1) | .013 (-5.0) |
| | | Right | -.00 (0.1) | -- | .001 (0.6) | .002 (0.9) | .000 (0.2) | .000 (0.4) | -.04 (11.9) | -0.011 | .000 (0.1) | .002 (1.3) |
| | **Frontal, pars orbitalis** | Left | -.03 (3.7) | -- | .000 (0.2) | .000 (0.3) | .003 (-1.5) | .004 (-1.8) | -.04 (13.7) | -0.020 | .001 (-0.8) | .011 (-4.3) |
| | | Right | -.02 (2.6) | -- | .000 (0.3) | .001 (0.6) | .002 (-0.9) | .006 (-2.4) | -.05 (18.6) | -0.008 | .001 (-0.5) | .017 (-6.5) |
| | **Frontal, lateral orbital** | Left | -.03 (2.0) | -- | .000 (-0.2) | .002 (-1.0) | .000 (0.2) | .003 (1.4) | -.05 (17.5) | 0.006 | .000 (0.3) | .034 (12.6) |
| | | Right | -.00 (0.1) | -- | .001 (0.5) | .001 (0.5) | .006 (2.3) | .008 (3.2) | -.04 (6.6) | -0.017 | .002 (-0.9) | .022 (-8.1) |
| | **Frontal, medial orbital** | Left | -.06 (7.4) | 0.103 | .000 (0.2) | .000 (0.1) | .000 (-0.4) | .000 (-0.0) | -.07 (17.4) | 0.010 | .000 (-0.2) | .003 (-1.5) |
| | | Right | -.06 (8.1) | 0.091 | .000 (-0.3) | .007 (-2.7) | .001 (0.7) | .001 (0.8) | -.04 (8.2) | -0.018 | .000 (-0.3) | .020 (-7.4) |
| | **Frontal, pole** | Left | -.03 (2.7) | -- | .000 (0.0) | .000 (0.4) | .000 (-0.2) | .000 (-0.4) | -.03 (6.1) | -0.018 | .000 (0.1) | .002 (1.1) |
| | | Right | -.03 (2.9) | -- | .000 (-0.4) | .005 (-2.0) | .001 (0.7) | .000 (0.0) | -.03 (3.7) | -- | .000 (0.2) | .003 (1.6) |
| | **Frontal, precentral gyrus** | Left | -.03 (3.2) | -- | .000 (-0.2) | .001 (-0.8) | .008 (-2.9) | .008 (-3.0) | -.06 (13.7) | 0.006 | .003 (-1.4) | .012 (-4.7) |
| | | Right | -.04 (4.4) | 0.112 | .000 (0.0) | .003 (1.2) | .001 (-0.7) | .001 (-0.8) | -.06 (12.2) | 0.015 | .002 (-1.0) | .019 (-7.1) |
| | **Parietal, postcentral gyrus** | Left | -.03 (3.6) | -- | .000 (0.1) | .000 (0.3) | .008 (-3.0) | .006 (-2.4) | -.05 (12.1) | 0.005 | .002 (-1.0) | .010 (-4.0) |
| | | Right | -.03 (2.6) | -- | .002 (0.9) | .001 (0.6) | .002 (-1.0) | .000 (-0.3) | -.05 (18.3) | 0.007 | .001 (-0.8) | .024 (-8.9) |
| | **Parietal, paracentral gyrus** | Left | -.05 (7.2) | 0.122 | .000 (0.0) | .001 (0.5) | .008 (-3.0) | .003 (-1.4) | -.05 (10.8) | -0.013 | .001 (-0.7) | .012 (-4.6) |
| | | Right | -.05 (9.2) | 0.109 | .000 (0.0) | .002 (0.9) | .000 (-0.3) | .000 (-0.0) | -.05 (8.0) | -0.020 | .001 (-0.8) | .021 (-7.8) |
| | **Parietal, superior** | Left | -.03 (4.3) | 0.121 | .000 (-0.1) | .001 (-0.7) | .003 (-1.3) | .002 (-1.0) | -.05 (11.2) | -0.003 | .000 (-0.3) | .019 (-7.1) |

| | | | | | | | | | | | | |
|---|---|---|---|---|---|---|---|---|---|---|---|---|
| | | Right | -.04 (4.7) | 0.101 | .000 (-0.3) | .006 (-2.3) | .001 (-0.8) | .001 (-0.5) | -.05 (11.8) | 0.002 | .000 (-0.4) | .011 (-4.1) |
| | **Parietal, inferior** | Left | -.04 (7.4) | 0.119 | .000 (0.0) | .001 (0.8) | .005 (-2.1) | .006 (-2.3) | -.05 (12.7) | -0.002 | .000 (-0.2) | .013 (-5.0) |
| | | Right | -.04 (6.9) | 0.097 | .001 (-0.4) | .006 (-2.3) | .000 (-0.1) | .001 (-0.5) | -.06 (18.9) | 0.007 | .001 (-0.6) | .014 (-5.3) |
| | **Parietal, supramarginal** | Left | -.04 (6.2) | 0.103 | .002 (-1.0) | .003 (-1.5) | .004 (-1.6) | .003 (-1.5) | -.05 (21.7) | 0.000 | .000 (-0.4) | .022 (-8.2) |
| | | Right | -.04 (5.5) | 0.096 | .000 (0.1) | .001 (0.7) | .000 (-0.1) | .000 (-0.3) | -.06 (24.8) | 0.006 | .000 (-0.3) | .020 (-7.6) |
| | **Parietal, precuneus** | Left | -.03 (5.7) | 0.102 | .000 (-0.2) | .002 (-0.9) | .004 (-1.8) | .004 (-1.6) | -.07 (29.2) | 0.003 | .000 (-0.3) | .011 (-4.2) |
| | | Right | -.03 (4.7) | 0.080 | .000 (-0.1) | .004 (-1.8) | .004 (-1.8) | .000 (-0.0) | -.06 (23.4) | 0.003 | .000 (-0.4) | .016 (-5.9) |
| | **Temporal, parahippocampal** | Left | -.03 (3.1) | -- | .000 (-0.2) | .000 (-0.1) | .001 (-0.6) | .000 (-0.2) | -.03 (11.7) | 0.015 | .003 (-1.4) | .023 (-8.4) |
| | | Right | -.01 (1.0) | -- | .000 (0.0) | .001 (0.4) | .000 (0.1) | .000 (0.3) | -.03 (9.4) | 0.014 | .000 (-0.1) | .029 (-10.5) |
| | **Temporal, entorhinal** | Left | -.06 (6.7) | 0.091 | .002 (-0.8) | .001 (-0.5) | .000 (-0.2) | .000 (-0.0) | -.05 (14.0) | 0.024 | .000 (-0.1) | .008 (-3.3) |
| | | Right | -.04 (4.7) | 0.067 | .000 (-0.3) | .002 (-1.0) | .000 (0.0) | .001 (0.7) | -.03 (11.1) | 0.025 | .002 (-1.2) | .033 (-11.8) |
| | **Temporal, pole** | Left | -- | -- | -- | -- | -- | -- | -.02 (3.7) | -- | .000 (0.0) | .002 (1.2) |
| | | Right | -- | -- | -- | -- | -- | -- | -.04 (8.4) | -0.001 | .000 (-0.4) | .009 (-3.8) |
| | **Temporal, superior** | Left | -.05 (9.7) | 0.125 | .000 (0.0) | .000 (0.2) | .007 (-2.9) | .004 (-1.8) | -.05 (39.0) | 0.004 | .001 (-0.5) | .031 (-11.5) |
| | | Right | -.06 (14.7) | 0.107 | .000 (-0.1) | .003 (-1.2) | .002 (-1.1) | .006 (-2.4) | -.06 (48.2) | 0.008 | .000 (-0.4) | .032 (-12.3) |
| | **Temporal, middle** | Left | -.06 (9.9) | 0.139 | .000 (0.1) | .000 (0.2) | .003 (-1.3) | .011 (-3.9) | -.05 (30.1) | 0.001 | .000 (-0.1) | .024 (-9.3) |
| | | Right | -.05 (5.7) | 0.122 | .000 (0.2) | .001 (0.8) | .000 (0.1) | .003 (1.4) | -.08 (46.9) | 0.016 | .001 (-0.5) | .032 (-12.3) |
| | **Temporal, inferior** | Left | -.05 (6.8) | 0.117 | .000 (0.2) | .000 (0.1) | .002 (-0.9) | .006 (-2.3) | -.06 (26.3) | 0.014 | .000 (0.2) | .033 (12.6) |
| | | Right | -.04 (4.6) | 0.099 | .000 (0.0) | .003 (1.3) | .000 (0.2) | .001 (0.7) | -.07 (37.2) | 0.019 | .000 (-0.4) | .024 (-9.1) |
| | **Temporal, transverse** | Left | -.01 (0.4) | -- | .000 (0.0) | .001 (0.7) | .003 (-1.5) | .000 (-0.0) | -.04 (7.4) | -0.007 | .000 (-0.4) | .000 (-0.1) |
| | | Right | -.02 (2.0) | 0.081 | .000 (-0.1) | .002 (-0.9) | .002 (-0.9) | .000 (-0.4) | -.02 (1.9) | -- | .000 (0.2) | .006 (2.6) |
| | **Temporal, bank sup temp sulc** | Left | -.03 (4.5) | 0.063 | .000 (-0.2) | .002 (-0.9) | .002 (-1.0) | .000 (-0.1) | -.05 (14.9) | 0.012 | .000 (0.0) | .008 (3.3) |
| | | Right | -.03 (5.1) | -- | .000 (-0.2) | .003 (-1.3) | .001 (-0.4) | .000 (-0.0) | -.07 (26.7) | 0.015 | .000 (-0.1) | .007 (-3.0) |
| | **Temporal, fusiform** | Left | -.07 (15.4) | 0.121 | .001 (-0.5) | .003 (-1.4) | .003 (-1.2) | .001 (-0.8) | -.07 (28.3) | 0.038 | .001 (-0.7) | .053 (-19.2) |

| | | | | | | | | | | | | |
|---|---|---|---|---|---|---|---|---|---|---|---|---|
| | | Right | -.03 (2.5) | -- | .000 (-0.2) | .003 (-1.3) | .000 (0.0) | .000 (0.1) | -.07 (30.6) | 0.035 | .001 (-0.8) | .026 (-9.7) |
| | **Occipital, lateral** | Left | -.04 (5.6) | 0.128 | .000 (0.3) | .000 (0.3) | .001 (-0.6) | .001 (-0.8) | -.05 (13.9) | 0.021 | .001 (-0.8) | .014 (-5.4) |
| | | Right | -.01 (0.9) | -- | .000 (0.3) | .002 (0.9) | .001 (-0.6) | .000 (-0.1) | -.06 (19.1) | 0.025 | .002 (-1.1) | .017 (-6.4) |
| | **Occipital, pericalcarine** | Left | -.00 (0.0) | -- | .001 (0.5) | .000 (0.1) | .001 (-0.4) | .002 (-0.8) | -.02 (2.3) | -- | .001 (-0.5) | .000 (-0.3) |
| | | Right | -.01 (0.7) | -- | .003 (1.3) | .001 (0.5) | .002 (1.0) | .004 (1.6) | -.04 (6.3) | 0.009 | .000 (-0.1) | .004 (-1.7) |
| | **Occipital, lingual** | Left | -.00 (0.1) | -- | .000 (0.2) | .000 (0.3) | .000 (-0.2) | .000 (-0.4) | -.05 (16.7) | 0.022 | .002 (-1.2) | .001 (-0.5) |
| | | Right | -.01 (0.4) | -- | .002 (0.9) | .001 (0.4) | .000 (0.2) | .003 (1.2) | -.06 (16.3) | 0.017 | .000 (-0.2) | .000 (-0.4) |
| | **Occipital, cuneus** | Left | -.01 (0.7) | -- | .000 (0.3) | .000 (0.1) | .002 (-0.9) | .000 (-0.3) | -.03 (7.9) | 0.007 | .000 (-0.2) | .000 (-0.1) |
| | | Right | -.01 (0.8) | -- | .001 (0.7) | .001 (0.4) | .001 (0.7) | .005 (2.1) | -.03 (5.0) | 0.003 | .000 (-0.1) | .001 (-0.4) |
| GWC | **Cingulate, caudal ant** | Left | -.06 (6.3) | 0.137 | .001 (0.7) | .001 (0.5) | .006 (-2.2) | .017 (-6.1) | -.04 (9.1) | 0.157 | .000 (-0.1) | .071 (-28.8) |
| | | Right | -.06 (7.8) | 0.131 | .000 (0.0) | .001 (0.5) | .003 (-1.5) | .010 (-3.6) | -.03 (5.3) | 0.144 | .001 (0.8) | .116 (46.4) |
| | **Cingulate, rostral ant** | Left | -.08 (21.2) | 0.080 | .002 (0.9) | .001 (0.6) | .017 (-5.9) | .032 (-10.9) | -.05 (9.2) | 0.135 | .001 (-0.4) | .064 (-24.6) |
| | | Right | -.07 (16.0) | 0.091 | .000 (0.0) | .000 (0.0) | .010 (-3.9) | .010 (-3.8) | -.04 (7.1) | 0.147 | .000 (0.4) | .085 (33.3) |
| | **Cingulate, posterior** | Left | -.10 (31.5) | 0.118 | .000 (-0.3) | .000 (-0.2) | .014 (-5.0) | .035 (-11.8) | -.04 (10.5) | 0.143 | .000 (0.4) | .079 (33.9) |
| | | Right | -.08 (22.5) | 0.104 | .000 (-0.1) | .000 (-0.4) | .013 (-4.6) | .035 (-12.0) | -.03 (5.0) | 0.148 | .000 (0.4) | .103 (42.5) |
| | **Cingulate, isthmus** | Left | -.01 (0.7) | -- | .001 (0.4) | .001 (0.5) | .003 (-1.3) | .005 (-2.2) | -.04 (8.5) | 0.132 | .000 (0.2) | .054 (23.1) |
| | | Right | -.02 (2.0) | -- | .001 (-0.6) | .001 (-0.7) | .001 (-0.7) | .001 (-0.5) | -.03 (4.2) | 0.134 | .000 (0.0) | .058 (24.5) |
| | **Insula** | Left | -.05 (10.2) | 0.076 | .000 (0.0) | .000 (0.1) | .000 (-0.2) | .002 (-0.9) | -.05 (16.0) | 0.149 | .001 (1.0) | .051 (22.2) |
| | | Right | -.02 (1.1) | -- | .000 (-0.4) | .001 (-0.4) | .004 (-1.8) | .007 (-2.7) | -.01 (1.6) | -- | .000 (0.1) | .042 (17.9) |
| | **Frontal, superior** | Left | -.13 (70.6) | 0.106 | .000 (0.0) | .000 (0.1) | .046 (-15.9) | .053 (-18.1) | -.05 (21.5) | 0.129 | .001 (0.8) | .147 (60.1) |
| | | Right | -.12 (67.5) | 0.104 | .000 (0.1) | .000 (0.1) | .046 (-15.5) | .057 (-19.4) | -.03 (7.8) | 0.127 | .000 (0.3) | .180 (72.0) |
| | **Frontal, caudal middle** | Left | -.11 (46.2) | 0.102 | .000 (0.1) | .000 (0.3) | .036 (-12.5) | .041 (-14.3) | -.04 (15.0) | 0.123 | .000 (0.4) | .117 (47.1) |
| | | Right | -.10 (39.3) | 0.096 | .000 (-0.1) | .000 (-0.0) | .038 (-12.8) | .048 (-16.1) | -.02 (4.1) | 0.120 | .000 (0.0) | .141 (54.4) |
| | **Frontal, rostral middle** | Left | -.10 (42.1) | 0.108 | .000 (0.2) | .000 (0.3) | .032 (-11.3) | .041 (-14.1) | -.06 (18.6) | 0.150 | .000 (0.5) | .150 (60.5) |

| | | | | | | | | | | | | |
|---|---|---|---|---|---|---|---|---|---|---|---|---|
| | | Right | -.11 (45.3) | 0.110 | .000 (0.2) | .000 (0.0) | .033 (-11.0) | .039 (-13.1) | -.04 (6.6) | 0.146 | .000 (0.1) | .175 (68.6) |
| | **Frontal, pars opercularis** | Left | -.08 (33.0) | 0.098 | .000 (-0.1) | .000 (-0.0) | .016 (-5.9) | .045 (-15.5) | -.05 (19.5) | 0.128 | .001 (0.9) | .098 (41.7) |
| | | Right | -.05 (13.3) | 0.084 | .000 (0.3) | .000 (0.2) | .021 (-7.3) | .041 (-13.6) | -.02 (3.4) | -- | .000 (0.0) | .122 (50.9) |
| | **Frontal, pars triangularis** | Left | -.07 (18.2) | 0.096 | .000 (-0.2) | .000 (-0.3) | .002 (-1.0) | .007 (-2.8) | -.05 (14.9) | 0.133 | .001 (0.7) | .100 (40.1) |
| | | Right | -.07 (18.0) | 0.090 | .000 (0.1) | .000 (0.2) | .008 (-3.0) | .014 (-5.0) | -.04 (7.0) | 0.144 | .000 (0.2) | .104 (40.8) |
| | **Frontal, pars orbitalis** | Left | -.08 (26.7) | 0.088 | .001 (0.6) | .001 (0.4) | .011 (-4.1) | .023 (-8.3) | -.05 (19.5) | 0.132 | .001 (0.7) | .107 (43.6) |
| | | Right | -.07 (23.7) | 0.089 | .000 (0.0) | .000 (0.1) | .021 (-7.3) | .030 (-10.1) | -.02 (4.3) | 0.131 | .000 (0.1) | .127 (51.7) |
| | **Frontal, lateral orbital** | Left | -.08 (25.9) | 0.085 | .000 (0.2) | .000 (0.1) | .001 (-0.8) | .005 (-2.2) | -.06 (22.6) | 0.140 | .001 (1.1) | .087 (36.6) |
| | | Right | -.06 (10.9) | 0.079 | .000 (0.0) | .000 (0.2) | .000 (0.0) | .001 (0.8) | -.03 (5.9) | 0.135 | .000 (-0.1) | .102 (-39.1) |
| | **Frontal, medial orbital** | Left | -.07 (15.8) | 0.078 | .000 (-0.3) | .000 (-0.1) | .012 (-4.3) | .019 (-6.8) | -.04 (10.0) | 0.129 | .000 (0.4) | .086 (32.9) |
| | | Right | -.08 (26.9) | 0.083 | .000 (0.1) | .000 (0.1) | .007 (-2.8) | .018 (-6.2) | -.04 (5.7) | 0.117 | .000 (0.1) | .078 (28.8) |
| | **Frontal, pole** | Left | -.10 (30.8) | 0.095 | .001 (0.5) | .000 (0.2) | .015 (-5.2) | .017 (-6.1) | -.07 (15.1) | 0.153 | .000 (0.4) | .094 (34.8) |
| | | Right | -.10 (30.9) | 0.094 | .000 (0.3) | .000 (0.1) | .009 (-3.3) | .016 (-5.8) | -.05 (9.1) | 0.140 | .000 (-0.1) | .095 (-35.1) |
| | **Frontal, precentral gyrus** | Left | -.10 (42.6) | 0.072 | .000 (0.0) | .001 (0.4) | .017 (-6.1) | .027 (-9.3) | -.04 (16.1) | 0.103 | .000 (0.3) | .063 (25.4) |
| | | Right | -.06 (19.8) | 0.064 | .000 (0.1) | .000 (0.3) | .025 (-8.6) | .046 (-15.7) | -.01 (1.4) | -- | .000 (0.2) | .075 (29.2) |
| | **Parietal, postcentral gyrus** | Left | -.09 (30.2) | 0.081 | .000 (-0.1) | .000 (-0.2) | .005 (-2.1) | .023 (-8.1) | -.04 (12.6) | 0.114 | .001 (0.8) | .034 (14.2) |
| | | Right | -.03 (4.6) | 0.060 | .001 (-0.5) | .000 (-0.4) | .009 (-3.3) | .031 (-10.5) | -.01 (0.8) | -- | .000 (0.1) | .039 (15.3) |
| | **Parietal, paracentral gyrus** | Left | -.09 (35.1) | 0.076 | .000 (-0.3) | .000 (-0.2) | .035 (-11.9) | .040 (-13.5) | -.04 (14.8) | 0.100 | .000 (0.3) | .053 (21.1) |
| | | Right | -.09 (29.9) | 0.070 | .000 (0.0) | .000 (0.2) | .022 (-7.8) | .036 (-12.5) | -.03 (8.4) | 0.104 | .000 (0.4) | .088 (34.2) |
| | **Parietal, superior** | Left | -.10 (40.2) | 0.105 | .000 (-0.2) | .000 (-0.1) | .024 (-8.4) | .037 (-12.8) | -.05 (20.6) | 0.130 | .000 (0.4) | .101 (42.2) |
| | | Right | -.05 (9.7) | 0.088 | .000 (-0.3) | .000 (-0.4) | .012 (-4.6) | .028 (-9.8) | -.04 (9.2) | 0.124 | .000 (0.1) | .092 (36.3) |
| | **Parietal, inferior** | Left | -.07 (23.0) | 0.098 | .000 (0.1) | .000 (0.0) | .011 (-4.2) | .022 (-7.7) | -.06 (24.5) | 0.137 | .001 (0.6) | .078 (33.1) |
| | | Right | -.02 (3.5) | -- | .000 (-0.4) | .000 (-0.3) | .003 (-1.4) | .003 (-1.4) | -.04 (10.9) | 0.123 | .000 (-0.2) | .089 (-37.2) |
| | **Parietal, supramarginal** | Left | -.09 (32.6) | 0.103 | .000 (-0.2) | .000 (-0.3) | .014 (-5.1) | .020 (-6.9) | -.05 (19.8) | 0.132 | .001 (0.7) | .086 (37.1) |

| | | | | | | | | | | | | |
|---|---|---|---|---|---|---|---|---|---|---|---|---|
| | | Right | -.01 (0.4) | -- | .000 (0.1) | .000 (0.2) | .021 (-7.3) | .023 (-8.2) | -.02 (3.3) | -- | .000 (-0.3) | .097 (-41.1) |
| | **Parietal, precuneus** | Left | -.07 (26.7) | 0.095 | .001 (-0.5) | .001 (-0.5) | .018 (-6.4) | .041 (-14.0) | -.05 (20.4) | 0.131 | .000 (0.3) | .096 (40.9) |
| | | Right | -.04 (8.5) | 0.081 | .000 (-0.2) | .001 (-0.4) | .012 (-4.7) | .030 (-10.6) | -.04 (14.7) | 0.125 | .000 (0.2) | .097 (41.1) |
| | **Temporal, parahippocampal** | Left | -.00 (0.1) | -- | .000 (-0.4) | .001 (-0.6) | .001 (0.8) | .007 (2.6) | -.04 (7.4) | 0.143 | .001 (0.7) | .023 (9.9) |
| | | Right | -.06 (11.1) | 0.062 | .000 (-0.2) | .001 (-0.5) | .002 (0.9) | .008 (3.0) | -.01 (1.1) | -- | .000 (0.2) | .014 (6.2) |
| | **Temporal, entorhinal** | Left | -.01 (0.6) | -- | .004 (-1.8) | .004 (-1.7) | .001 (0.6) | .000 (0.2) | -.05 (11.5) | 0.116 | .002 (1.1) | .010 (4.4) |
| | | Right | -.01 (1.0) | -- | .002 (-1.2) | .003 (-1.6) | .000 (0.1) | .002 (1.2) | -.02 (2.6) | -- | .001 (0.9) | .034 (13.5) |
| | **Temporal, pole** | Left | -.03 (2.4) | -- | .000 (0.1) | .000 (0.1) | .002 (-1.0) | .006 (-2.4) | -.04 (4.6) | 0.146 | .000 (0.5) | .035 (13.6) |
| | | Right | -.01 (0.6) | -- | .001 (-0.8) | .002 (-1.0) | .000 (0.0) | .000 (0.1) | -.04 (7.9) | 0.114 | .000 (0.0) | .031 (12.3) |
| | **Temporal, superior** | Left | -.05 (10.8) | 0.093 | .000 (0.0) | .000 (0.0) | .005 (-2.1) | .016 (-5.7) | -.05 (20.4) | 0.140 | .000 (0.4) | .070 (32.4) |
| | | Right | -.03 (4.5) | 0.065 | .001 (-0.5) | .001 (-0.7) | .009 (-3.4) | .029 (-10.0) | -.01 (1.1) | -- | .000 (0.0) | .075 (34.6) |
| | **Temporal, middle** | Left | -.03 (4.6) | 0.089 | .000 (0.2) | .000 (0.0) | .002 (-1.2) | .012 (-4.3) | -.06 (26.3) | 0.136 | .000 (0.2) | .064 (30.3) |
| | | Right | -.07 (22.4) | 0.053 | .001 (-0.7) | .002 (-0.9) | .001 (-0.7) | .011 (-4.2) | -.01 (0.9) | -- | .000 (-0.4) | .079 (-36.7) |
| | **Temporal, inferior** | Left | -.00 (0.2) | -- | .000 (0.0) | .000 (0.2) | .000 (0.0) | .008 (3.0) | -.06 (23.9) | 0.140 | .000 (0.5) | .060 (28.1) |
| | | Right | -.07 (26.5) | 0.041 | .003 (-1.2) | .006 (-2.5) | .000 (0.0) | .004 (1.7) | -.02 (2.3) | -- | .000 (-0.2) | .054 (-25.0) |
| | **Temporal, transverse** | Left | -.02 (2.2) | -- | .000 (0.0) | .000 (0.1) | .000 (-0.1) | .004 (-1.9) | -.05 (16.4) | 0.108 | .000 (0.5) | .044 (17.7) |
| | | Right | -.04 (6.7) | 0.029 | .000 (0.0) | .000 (0.3) | .003 (-1.3) | .007 (-2.6) | -.02 (4.7) | 0.091 | .000 (0.0) | .045 (17.9) |
| | **Temporal, bank sup temp sulc** | Left | -.01 (1.1) | -- | .000 (0.1) | .000 (0.0) | .003 (-1.3) | .006 (-2.5) | -.05 (15.0) | 0.132 | .000 (0.3) | .047 (20.7) |
| | | Right | -.07 (24.7) | 0.047 | .000 (-0.1) | .000 (-0.0) | .002 (-0.9) | .003 (-1.5) | -.01 (0.5) | -- | .000 (-0.1) | .055 (-23.8) |
| | **Temporal, fusiform** | Left | -.00 (0.0) | -- | .000 (-0.1) | .001 (-0.7) | .000 (0.0) | .007 (2.7) | -.06 (24.7) | 0.134 | .000 (0.4) | .055 (25.9) |
| | | Right | -.06 (24.1) | 0.036 | .001 (-0.5) | .003 (-1.5) | .000 (0.4) | .001 (0.8) | -.03 (5.8) | 0.117 | .000 (0.1) | .060 (29.6) |
| | **Occipital, lateral** | Left | -.02 (1.9) | -- | .000 (0.2) | .000 (0.1) | .001 (0.5) | .003 (1.5) | -.06 (16.2) | 0.141 | .000 (0.3) | .034 (15.0) |
| | | Right | -.12 (51.5) | 0.025 | .000 (-0.1) | .001 (-0.7) | .004 (1.7) | .001 (0.7) | -.02 (3.1) | -- | .000 (-0.3) | .059 (-25.7) |
| | **Occipital, pericalcarine** | Left | -.02 (2.5) | -- | .003 (-1.3) | .001 (-0.5) | .008 (2.9) | .000 (0.1) | -.03 (3.0) | -- | .000 (0.3) | .006 (2.6) |

| | | | | | | | | | | | | |
|---|---|---|---|---|---|---|---|---|---|---|---|---|
| | | Right | -.06 (13.7) | 0.016 | .000 (-0.4) | .001 (-0.5) | .018 (6.3) | .003 (1.5) | -.00 (0.1) | -- | .000 (0.2) | .011 (4.6) |
| | **Occipital, lingual** | Left | -.04 (9.0) | 0.034 | .001 (-0.5) | .001 (-0.4) | .001 (0.5) | .000 (0.0) | -.04 (6.0) | 0.135 | .001 (0.7) | .022 (9.6) |
| | | Right | -.08 (26.1) | 0.036 | .000 (-0.3) | .004 (-1.7) | .014 (5.1) | .006 (2.6) | -.01 (0.9) | -- | .000 (0.0) | .031 (13.2) |
| | **Occipital, cuneus** | Left | -.01 (0.5) | -- | .000 (0.3) | .000 (0.1) | .001 (0.8) | .000 (0.4) | -.03 (4.9) | 0.144 | .001 (0.7) | .050 (20.4) |
| | | Right | -.05 (11.5) | 0.037 | .000 (0.1) | .001 (0.6) | .011 (4.1) | .000 (0.3) | -.02 (1.6) | -- | .000 (0.0) | .059 (23.8) |
| Volume (cortical) | **Cingulate, caudal ant** | Left | -.02 (2.5) | -- | .002 (1.2) | .000 (0.3) | .000 (-0.4) | .001 (-0.8) | -.02 (33.9) | 0.000 | .000 (-0.1) | .015 (-5.6) |
| | | Right | -.02 (2.9) | -- | .000 (0.0) | .002 (0.9) | .001 (-0.7) | .002 (-1.2) | -.02 (32.2) | -0.005 | .002 (-1.4) | .006 (-2.7) |
| | **Cingulate, rostral ant** | Left | -.03 (7.5) | 0.050 | .001 (-0.5) | .002 (-0.9) | .001 (-0.6) | .003 (-1.5) | -.03 (32.3) | 0.001 | .002 (0.9) | .018 (6.8) |
| | | Right | -.02 (7.4) | 0.032 | .000 (0.3) | .001 (0.5) | .007 (-2.8) | .002 (-0.9) | -.02 (20.4) | -0.006 | .000 (-0.4) | .002 (-1.3) |
| | **Cingulate, posterior** | Left | -.03 (9.3) | 0.062 | .000 (-0.3) | .003 (-1.2) | .001 (0.5) | .001 (0.5) | -.04 (59.2) | 0.001 | .000 (-0.1) | .021 (-7.9) |
| | | Right | -.03 (5.7) | 0.027 | .000 (-0.1) | .000 (-0.4) | .006 (-2.4) | .003 (-1.3) | -.03 (47.7) | -0.002 | .000 (-0.2) | .003 (-1.3) |
| | **Cingulate, isthmus** | Left | -.02 (6.8) | 0.038 | .000 (0.1) | .000 (0.4) | .001 (-0.5) | .000 (-0.3) | -.03 (40.4) | 0.004 | .000 (-0.2) | .017 (-6.2) |
| | | Right | -.02 (6.1) | 0.039 | .000 (-0.3) | .005 (-1.9) | .001 (0.7) | .003 (1.5) | -.03 (30.7) | 0.009 | .000 (0.0) | .006 (2.7) |
| | **Insula** | Left | -.02 (3.4) | -- | .000 (-0.2) | .001 (-0.6) | .003 (-1.3) | .001 (-0.8) | -.03 (29.0) | 0.000 | .000 (-0.4) | .003 (-1.2) |
| | | Right | -.02 (3.0) | -- | .000 (-0.1) | .000 (-0.2) | .000 (0.0) | .000 (0.2) | -.03 (25.3) | 0.007 | .002 (-0.9) | .009 (-3.6) |
| | **Frontal, superior** | Left | -.03 (12.4) | 0.065 | .000 (0.1) | .001 (0.5) | .002 (-1.1) | .009 (-3.3) | -.04 (38.9) | -0.015 | .000 (-0.1) | .012 (-4.6) |
| | | Right | -.03 (8.0) | 0.055 | .000 (0.3) | .003 (1.6) | .000 (-0.1) | .002 (-1.0) | -.03 (32.6) | -0.015 | .000 (-0.4) | .010 (-4.0) |
| | **Frontal, caudal middle** | Left | -.01 (2.8) | -- | .001 (0.7) | .000 (0.0) | .000 (-0.3) | .002 (-1.0) | -.03 (30.5) | -0.009 | .000 (-0.5) | .009 (-3.4) |
| | | Right | -.02 (3.5) | -- | .000 (0.3) | .002 (1.1) | .000 (-0.1) | .000 (-0.2) | -.03 (23.9) | -0.012 | .000 (-0.5) | .006 (-2.6) |
| | **Frontal, rostral middle** | Left | -.04 (20.1) | 0.061 | .000 (0.2) | .001 (0.8) | .002 (-1.1) | .016 (-5.7) | -.04 (50.5) | -0.003 | .000 (-0.5) | .031 (-11.2) |
| | | Right | -.03 (11.8) | 0.049 | .001 (0.4) | .004 (1.7) | .002 (-1.1) | .008 (-3.0) | -.03 (45.5) | -0.004 | .000 (0.0) | .033 (12.2) |
| | **Frontal, pars opercularis** | Left | -.03 (11.5) | 0.052 | .001 (0.5) | .000 (0.2) | .002 (-0.8) | .009 (-3.2) | -.03 (47.1) | -0.003 | .002 (-1.1) | .011 (-4.4) |
| | | Right | -.03 (8.4) | 0.037 | .001 (0.7) | .001 (0.6) | .000 (-0.1) | .000 (-0.0) | -.03 (43.1) | -0.003 | .000 (-0.5) | .017 (-6.5) |
| | **Frontal, pars triangularis** | Left | -.05 (15.3) | 0.050 | .000 (-0.1) | .002 (-0.9) | .001 (-0.7) | .010 (-3.8) | -.04 (33.9) | -0.006 | .000 (-0.2) | .009 (-3.8) |

| | | | | | | | | | | | | |
|---|---|---|---|---|---|---|---|---|---|---|---|---|
| | | Right | -.03 (5.1) | 0.039 | .001 (0.7) | .002 (0.9) | .000 (-0.3) | .003 (-1.5) | -.04 (48.9) | 0.001 | .000 (0.3) | .017 (6.3) |
| | **Frontal, pars orbitalis** | Left | -.04 (19.2) | 0.069 | .000 (0.0) | .001 (0.6) | .000 (-0.4) | .006 (-2.3) | -.03 (44.4) | -0.006 | .000 (-0.2) | .013 (-5.1) |
| | | Right | -.03 (9.0) | 0.055 | .001 (0.6) | .000 (0.3) | .001 (-0.7) | .007 (-2.7) | -.03 (54.9) | 0.000 | .000 (-0.4) | .023 (-8.7) |
| | **Frontal, lateral orbital** | Left | -.04 (15.6) | 0.066 | .000 (-0.1) | .002 (-1.0) | .002 (-0.9) | .002 (-1.0) | -.04 (44.8) | 0.012 | .000 (0.2) | .058 (20.7) |
| | | Right | -.02 (2.7) | -- | .000 (0.1) | .003 (1.3) | .000 (-0.3) | .000 (-0.3) | -.03 (16.2) | 0.008 | .000 (-0.4) | .022 (-8.2) |
| | **Frontal, medial orbital** | Left | -.05 (14.4) | 0.075 | .000 (0.1) | .002 (0.9) | .000 (0.2) | .000 (0.2) | -.04 (28.8) | 0.015 | .000 (-0.1) | .013 (-5.0) |
| | | Right | -.05 (12.9) | 0.061 | .000 (-0.3) | .006 (-2.2) | .001 (0.5) | .000 (0.0) | -.03 (14.6) | 0.013 | .002 (-1.1) | .043 (-15.2) |
| | **Frontal, pole** | Left | -.04 (5.2) | 0.067 | .000 (0.0) | .002 (0.9) | .000 (0.0) | .001 (0.5) | -.06 (41.2) | -0.001 | .000 (0.2) | .012 (5.1) |
| | | Right | -.03 (2.7) | -- | .000 (-0.1) | .002 (-1.1) | .002 (1.2) | .001 (0.7) | -.05 (28.3) | 0.000 | .000 (0.3) | .012 (5.0) |
| | **Frontal, precentral gyrus** | Left | -.03 (7.5) | 0.078 | .000 (0.0) | .001 (0.5) | .010 (-3.7) | .008 (-3.1) | -.04 (25.1) | -0.007 | .001 (-0.9) | .009 (-3.6) |
| | | Right | -.03 (6.4) | 0.073 | .000 (0.4) | .002 (0.8) | .003 (-1.4) | .001 (-0.6) | -.04 (24.1) | -0.004 | .002 (-1.0) | .010 (-4.2) |
| | **Parietal, postcentral gyrus** | Left | -.03 (8.0) | 0.069 | .000 (0.1) | .000 (0.3) | .008 (-3.1) | .005 (-2.1) | -.03 (27.0) | -0.004 | .000 (-0.4) | .008 (-3.3) |
| | | Right | -.03 (5.2) | 0.065 | .001 (0.7) | .001 (0.7) | .001 (-0.6) | .001 (-0.7) | -.04 (30.1) | -0.003 | .001 (-0.5) | .017 (-6.2) |
| | **Parietal, paracentral gyrus** | Left | -.03 (8.1) | 0.077 | .000 (0.2) | .000 (0.2) | .008 (-3.0) | .002 (-1.2) | -.04 (27.2) | -0.005 | .000 (-0.3) | .021 (-7.9) |
| | | Right | -.03 (8.2) | 0.058 | .000 (0.0) | .001 (0.7) | .002 (-1.1) | .000 (-0.0) | -.03 (16.9) | -0.016 | .001 (-0.5) | .015 (-5.8) |
| | **Parietal, superior** | Left | -.03 (9.6) | 0.072 | .000 (-0.2) | .002 (-1.0) | .002 (-0.9) | .005 (-2.0) | -.04 (37.3) | -0.002 | .000 (-0.4) | .020 (-7.6) |
| | | Right | -.03 (7.9) | 0.064 | .000 (0.0) | .003 (1.5) | .000 (-0.2) | .001 (-0.8) | -.04 (37.3) | -0.003 | .001 (-0.5) | .010 (-3.9) |
| | **Parietal, inferior** | Left | -.04 (25.5) | 0.063 | .000 (0.1) | .002 (1.0) | .002 (-1.2) | .010 (-3.8) | -.04 (60.4) | 0.004 | .000 (0.1) | .026 (9.4) |
| | | Right | -.04 (22.8) | 0.050 | .000 (0.1) | .002 (0.9) | .000 (-0.1) | .002 (-0.9) | -.04 (72.0) | 0.005 | .000 (-0.4) | .014 (-5.4) |
| | **Parietal, supramarginal** | Left | -.03 (14.0) | 0.052 | .001 (-0.5) | .004 (-1.7) | .001 (-0.8) | .007 (-2.6) | -.03 (54.3) | 0.002 | .000 (0.0) | .024 (8.9) |
| | | Right | -.03 (9.3) | 0.044 | .000 (0.1) | .001 (0.5) | .000 (0.1) | .001 (0.4) | -.04 (58.8) | 0.003 | .000 (-0.1) | .018 (-6.6) |
| | **Parietal, precuneus** | Left | -.04 (15.9) | 0.067 | .000 (0.1) | .002 (1.0) | .000 (-0.4) | .005 (-2.1) | -.04 (62.4) | 0.006 | .000 (0.0) | .017 (6.2) |
| | | Right | -.03 (11.4) | 0.057 | .001 (0.8) | .001 (0.8) | .000 (0.0) | .000 (0.3) | -.04 (55.8) | 0.004 | .000 (-0.4) | .015 (-5.7) |

| | | | | | | | | | | | | |
|---|---|---|---|---|---|---|---|---|---|---|---|---|
| | **Temporal, parahippocampal** | Left | -.04 (8.5) | 0.070 | .001 (-0.5) | .001 (-0.5) | .002 (-1.0) | .000 (-0.2) | -.03 (28.3) | 0.014 | .000 (-0.3) | .007 (-2.8) |
| | | Right | -.03 (4.5) | 0.056 | .000 (0.1) | .001 (0.5) | .000 (0.2) | .001 (0.5) | -.03 (23.0) | 0.017 | .001 (-0.8) | .048 (-17.0) |
| | **Temporal, entorhinal** | Left | -.03 (2.9) | -- | .000 (-0.1) | .000 (-0.1) | .001 (0.6) | .000 (0.2) | -.03 (13.1) | 0.009 | .000 (0.0) | .009 (3.7) |
| | | Right | -.01 (1.1) | -- | .002 (0.8) | .000 (0.2) | .001 (0.4) | .002 (0.8) | -.02 (7.4) | 0.009 | .001 (-0.6) | .000 (-0.1) |
| | **Temporal, pole** | Left | -- | -- | -- | -- | -- | -- | -.04 (12.4) | 0.014 | .001 (0.6) | .009 (3.6) |
| | | Right | -- | -- | -- | -- | -- | -- | -.05 (28.8) | 0.015 | .001 (-0.7) | .013 (-5.2) |
| | **Temporal, superior** | Left | -.04 (20.3) | 0.068 | .000 (0.4) | .000 (0.3) | .004 (-1.8) | .007 (-2.8) | -.04 (96.9) | 0.007 | .000 (0.0) | .030 (11.1) |
| | | Right | -.05 (28.6) | 0.053 | .000 (0.1) | .003 (1.2) | .001 (-0.6) | .002 (-1.2) | -.04 (103.7) | 0.008 | .001 (-0.8) | .030 (-10.9) |
| | **Temporal, middle** | Left | -.06 (35.6) | 0.070 | .000 (0.2) | .001 (0.5) | .003 (-1.2) | .011 (-4.1) | -.04 (108.2) | 0.014 | .001 (0.5) | .040 (14.1) |
| | | Right | -.04 (22.6) | 0.057 | .000 (0.3) | .003 (1.4) | .001 (-0.7) | .007 (-2.6) | -.05 (127.8) | 0.014 | .000 (0.0) | .042 (15.3) |
| | **Temporal, inferior** | Left | -.05 (29.2) | 0.054 | .001 (0.7) | .000 (0.0) | .003 (-1.5) | .008 (-3.2) | -.04 (69.3) | 0.019 | .001 (0.4) | .048 (17.2) |
| | | Right | -.03 (16.4) | 0.043 | .001 (0.8) | .000 (0.3) | .000 (-0.2) | .003 (-1.3) | -.04 (84.5) | 0.019 | .000 (0.0) | .046 (16.2) |
| | **Temporal, transverse** | Left | -.02 (3.8) | -- | .000 (0.3) | .000 (0.3) | .000 (-0.1) | .000 (-0.1) | -.04 (42.3) | 0.002 | .000 (-0.2) | .001 (-0.9) |
| | | Right | -.02 (4.6) | 0.038 | .001 (-0.7) | .004 (-1.8) | .001 (-0.4) | .000 (-0.1) | -.03 (24.4) | 0.012 | .000 (0.2) | .010 (4.1) |
| | **Temporal, bank sup temp sulc** | Left | -.04 (20.0) | 0.036 | .000 (0.0) | .001 (0.6) | .002 (-0.9) | .001 (-0.5) | -.02 (28.8) | 0.006 | .000 (-0.3) | .014 (-5.4) |
| | | Right | -.04 (15.4) | 0.023 | .000 (-0.1) | .000 (-0.3) | .002 (-1.1) | .001 (-0.7) | -.04 (48.3) | 0.008 | .000 (-0.4) | .006 (-2.8) |
| | **Temporal, fusiform** | Left | -.05 (27.9) | 0.060 | .000 (0.0) | .000 (0.3) | .002 (-1.1) | .003 (-1.4) | -.04 (66.1) | 0.023 | .000 (-0.2) | .044 (-15.9) |
| | | Right | -.03 (11.6) | 0.055 | .002 (0.8) | .000 (0.3) | .000 (0.0) | .001 (0.5) | -.04 (68.4) | 0.021 | .001 (-0.4) | .026 (-9.6) |
| | **Occipital, lateral** | Left | -.04 (14.8) | 0.078 | .000 (0.3) | .000 (0.4) | .000 (-0.4) | .004 (-1.8) | -.05 (87.0) | 0.021 | .000 (-0.1) | .017 (-6.3) |
| | | Right | -.03 (7.1) | 0.049 | .001 (0.7) | .002 (0.8) | .000 (-0.3) | .000 (-0.2) | -.05 (78.1) | 0.022 | .001 (-0.5) | .029 (-10.6) |
| | **Occipital, pericalcarine** | Left | -.00 (0.4) | -- | .005 (2.1) | .000 (0.2) | .000 (0.0) | .000 (0.0) | -.02 (5.5) | 0.011 | .000 (-0.2) | .001 (-0.7) |
| | | Right | -.01 (2.3) | -- | .006 (2.3) | .000 (0.0) | .002 (1.2) | .004 (1.9) | -.02 (8.5) | 0.010 | .000 (0.1) | .005 (2.3) |
| | **Occipital, lingual** | Left | -.02 (3.7) | -- | .001 (0.5) | .000 (0.1) | .003 (-1.4) | .000 (-0.2) | -.04 (61.3) | 0.016 | .000 (-0.3) | .010 (-4.1) |
| | | Right | -.02 (2.5) | -- | .002 (0.8) | .001 (0.6) | .000 (0.3) | .001 (0.6) | -.04 (51.3) | 0.017 | .000 (-0.2) | .007 (-2.9) |

| | | | | | | | | | | | | |
|---|---|---|---|---|---|---|---|---|---|---|---|---|
| | **Occipital, cuneus** | Left | -.02 (5.0) | 0.062 | .001 (0.4) | .000 (0.3) | .000 (-0.2) | .000 (-0.1) | -.03 (31.8) | 0.010 | .000 (0.2) | .006 (2.5) |
| | | Right | -.02 (4.2) | 0.049 | .001 (0.5) | .001 (0.6) | .002 (1.2) | .002 (0.8) | -.03 (24.2) | 0.010 | .001 (-0.6) | .005 (-2.3) |
| Volume (subc.) | **3rd Ventricle** | Bil | .06 (67.9) | -0.019 | .001 (0.4) | .000 (0.2) | .001 (0.6) | .003 (1.4) | -.05 (123.5) | -0.004 | .008 (3.4) | .002 (1.2) |
| | **4th Ventricle** | Bil | .02 (11.4) | 0.000 | .001 (0.4) | .001 (0.5) | .011 (4.1) | .004 (1.8) | -.02 (17.6) | 0.015 | .008 (3.2) | .007 (2.8) |
| | **5th Ventricle** | Bil | .00 (0.1) | -- | .000 (0.0) | .000 (0.1) | .001 (-0.6) | .000 (-0.3) | -.05 (3.4) | -- | .002 (-0.9) | .005 (-2.1) |
| | **Inf lat vent** | Left | .05 (38.3) | -0.002 | .007 (3.0) | .001 (0.7) | .013 (4.9) | .006 (2.4) | -.06 (71.1) | 0.002 | .024 (9.3) | .009 (5.0) |
| | | Right | .05 (44.0) | -0.008 | .031 (11.2) | .018 (6.7) | .011 (4.2) | .009 (3.6) | -.06 (62.8) | 0.002 | .020 (7.6) | .005 (2.7) |
| | **Lat vent** | Left | .05 (121.2) | -0.005 | .032 (12.2) | .016 (6.4) | .013 (5.2) | .007 (3.0) | -.05 (171.2) | -0.008 | .025 (10.9) | .012 (6.2) |
| | | Right | .05 (123.9) | -0.005 | .031 (11.7) | .015 (6.0) | .010 (4.1) | .005 (2.3) | -.05 (182.8) | -0.009 | .026 (11.7) | .011 (6.2) |
| | **CSF total** | Bil | -.03 (8.1) | -0.046 | .001 (-0.8) | .001 (-0.4) | .010 (-3.7) | .001 (-0.8) | -.03 (17.3) | -0.009 | .010 (3.6) | .003 (1.5) |
| | **Accumbens** | Left | -.08 (26.7) | 0.038 | .000 (0.4) | .000 (0.3) | .003 (-1.3) | .005 (-2.0) | -.04 (9.4) | 0.033 | .002 (-1.0) | .017 (-6.4) |
| | | Right | -.05 (16.7) | 0.034 | .000 (-0.3) | .001 (-0.8) | .003 (-1.5) | .000 (-0.0) | -.02 (5.7) | -0.003 | .000 (0.0) | .007 (2.9) |
| | **Amygdala** | Left | -.05 (15.3) | 0.030 | .000 (-0.3) | .001 (-0.6) | .008 (-3.1) | .003 (-1.3) | -.03 (13.5) | 0.007 | .000 (0.3) | .006 (2.6) |
| | | Right | -.04 (11.5) | 0.011 | .000 (0.4) | .000 (0.4) | .002 (-1.1) | .000 (-0.1) | -.03 (15.4) | 0.003 | .003 (-1.5) | .002 (-1.4) |
| | **Brainstem** | Bil | -.02 (10.2) | 0.012 | .002 (0.9) | .000 (0.0) | .001 (0.8) | .003 (1.2) | -.03 (26.7) | 0.013 | .001 (-0.4) | .043 (-15.8) |
| | **Caudate** | Left | -.02 (5.7) | 0.027 | .015 (5.2) | .004 (1.7) | .000 (0.0) | .001 (0.6) | -.03 (20.9) | -0.003 | .000 (-0.1) | .000 (-0.1) |
| | | Right | -.01 (1.6) | -- | .011 (4.0) | .003 (1.4) | .000 (0.3) | .017 (6.0) | -.03 (37.5) | 0.004 | .000 (-0.4) | .004 (-1.8) |
| | **Cerebellum** | Left | -.02 (3.8) | -- | .002 (1.2) | .000 (0.1) | .002 (1.2) | .001 (0.4) | -.04 (44.2) | 0.004 | .000 (-0.4) | .005 (-2.2) |
| | | Right | -.02 (6.5) | 0.019 | .003 (1.5) | .001 (0.6) | .000 (0.3) | .000 (0.3) | -.04 (30.9) | 0.006 | .000 (0.2) | .005 (2.4) |
| | **Hippocampus** | Left | -.06 (28.6) | 0.030 | .002 (1.1) | .001 (0.7) | .003 (-1.3) | .002 (-1.0) | -.06 (81.6) | -0.005 | .001 (-0.6) | .000 (-0.1) |
| | | Right | -.06 (46.7) | 0.019 | .000 (-0.1) | .000 (-0.1) | .001 (-0.5) | .000 (-0.0) | -.05 (69.6) | -0.010 | .000 (-0.2) | .002 (-1.2) |
| | **Pallidum** | Left | -.02 (1.9) | -- | .000 (0.3) | .001 (0.5) | .001 (0.7) | .002 (0.9) | -.01 (1.4) | -- | .001 (0.8) | .045 (16.0) |
| | | Right | -.01 (1.7) | -- | .003 (-1.5) | .003 (-1.4) | .000 (-0.2) | .004 (-1.6) | -.00 (0.2) | -- | .000 (-0.2) | .017 (-6.4) |
| | **Putamen** | Left | -.05 (22.6) | 0.031 | .005 (2.0) | .001 (0.6) | .003 (-1.3) | .009 (-3.6) | -.03 (23.1) | 0.005 | .000 (-0.3) | .006 (-2.7) |

| | | | | | | | | | | | | |
|---|---|---|---|---|---|---|---|---|---|---|---|---|
| | | Right | -.04 (18.8) | 0.020 | .006 (2.6) | .002 (0.9) | .004 (-1.7) | .003 (-1.5) | -.04 (56.8) | 0.003 | .000 (-0.2) | .001 (-0.4) |
| | **Thalamus** | Left | -.07 (25.2) | 0.010 | .000 (-0.1) | .000 (-0.0) | .003 (-1.3) | .009 (-3.5) | -.04 (40.2) | -0.006 | .002 (-1.0) | .037 (-13.9) |
| | | Right | -.07 (32.8) | 0.007 | .000 (0.0) | .000 (0.1) | .003 (-1.3) | .004 (-1.8) | -.06 (69.0) | -0.011 | .001 (-0.5) | .002 (-1.4) |
| | **CC anterior** | Bil | -.01 (0.7) | -- | .000 (-0.2) | .000 (-0.0) | .001 (0.8) | .000 (0.1) | -.02 (6.5) | 0.000 | .002 (-1.1) | .000 (-0.2) |
| | **CC central** | Bil | -.04 (9.0) | 0.005 | .000 (0.2) | .000 (0.1) | .001 (-0.7) | .000 (-0.1) | -.02 (9.6) | -0.004 | .000 (-0.1) | .001 (-0.8) |
| | **CC mid anterior** | Bil | -.04 (8.6) | -0.001 | .001 (0.6) | .000 (0.3) | .000 (-0.2) | .001 (-0.7) | -.03 (8.8) | -0.004 | .001 (0.7) | .003 (1.5) |
| | **CC mid posterior** | Bil | -.03 (7.8) | 0.012 | .000 (-0.1) | .000 (-0.0) | .000 (-0.3) | .000 (-0.0) | -.03 (16.0) | -0.007 | .005 (-2.0) | .002 (-1.2) |
| | **CC posterior** | Bil | -.01 (0.7) | -- | .000 (0.3) | .000 (0.3) | .000 (0.4) | .001 (0.5) | -.01 (1.4) | -- | .000 (-0.3) | .001 (-0.8) |
| | **Cerebellum WM** | Left | -.02 (2.5) | -- | .000 (0.2) | .001 (0.4) | .000 (0.0) | .004 (1.6) | -.02 (5.2) | -0.009 | .000 (0.4) | .000 (0.1) |
| | | Right | -.03 (3.2) | -- | .001 (-0.7) | .002 (-0.9) | .000 (0.2) | .008 (3.0) | -.03 (7.0) | -0.007 | .000 (0.0) | .001 (0.6) |
| | **Cerebral WM** | Left | -.03 (45.5) | 0.001 | .001 (-0.6) | .001 (-0.8) | .006 (2.5) | .002 (1.1) | -.04 (56.2) | 0.001 | .000 (-0.2) | .000 (-0.4) |
| | | Right | -.03 (44.2) | -0.005 | .003 (-1.2) | .002 (-1.0) | .005 (2.3) | .001 (0.5) | -.03 (38.5) | -0.002 | .001 (-0.6) | .000 (-0.1) |
| | | Bil | -- | -- | -- | -- | -- | -- | -.03 (53.1) | 0.000 | .001 (-0.4) | .000 (-0.1) |
| | **WM hypointensities** | Bil | .06 (35.0) | -0.031 | .021 (7.7) | .024 (8.8) | .000 (-0.1) | .008 (-3.3) | -.04 (56.0) | -0.009 | .020 (7.9) | .001 (0.7) |
| | **Non WM hypointensities** | Bil | .00 (0.3) | -- | .002 (1.0) | .001 (0.7) | .000 (0.1) | .001 (0.6) | -.03 (3.6) | -- | .000 (0.1) | .002 (1.0) |
| | **Brain Seg** | Bil | -.03 (31.9) | 0.036 | .001 (0.4) | .001 (0.6) | .000 (0.0) | .005 (2.0) | -.04 (129.5) | 0.001 | .000 (-0.1) | .022 (-8.2) |
| | **Brain Seg not vent** | Bil | -0.04 (49.2) | 0.039 | .000 (0.1) | .002 (1.0) | .000 (-0.2) | .006 (-2.6) | -.04 (160.2) | 0.002 | .001 (-0.7) | .024 (-9.3) |
| | **Brain Seg not vent surf** | Bil | -.04 (58.3) | 0.038 | .000 (0.2) | .003 (1.5) | .000 (0.1) | .008 (3.2) | -.05 (161.3) | 0.002 | .001 (-0.7) | .023 (-9.1) |
| | **Cortex** | Left | -.05 (21.0) | 0.091 | .000 (0.2) | .001 (0.6) | .003 (-1.5) | .008 (-3.2) | -.05 (100.6) | 0.006 | .000 (-0.2) | .045 (-16.3) |
| | | Right | -.04 (15.0) | 0.074 | .001 (0.6) | .003 (1.3) | .000 (-0.3) | .001 (-0.8) | -.05 (96.3) | 0.005 | .001 (-0.6) | .041 (-14.7) |
| | | Bil | -- | -- | -- | -- | -- | -- | -.05 (105.7) | 0.005 | .000 (-0.4) | .047 (-16.8) |
| | **eICV** | Bil | -.00 (0.8) | -- | .001 (0.5) | .000 (0.3) | .000 (0.3) | .008 (3.1) | -.00 (0.5) | -- | .002 (1.3) | .000 (0.0) |
| | **Subcort gray** | Bil | -.06 (59.7) | 0.021 | .003 (1.3) | .000 (0.3) | .003 (-1.5) | .003 (-1.5) | -.05 (113.5) | -0.005 | .002 (-1.1) | .001 (-0.9) |
| | **Supratentorial** | Bil | -.03 (42.4) | 0.036 | .001 (0.4) | .002 (1.1) | .000 (0.1) | .006 (2.3) | -.04 (113.2) | 0.001 | .000 (-0.1) | .018 (-7.0) |

| | | | | | | | | | | | | |
|---|---|---|---|---|---|---|---|---|---|---|---|---|
| | **Supratentorial not vent** | Bil | -.04 (66.9) | 0.038 | .000 (0.0) | .004 (1.9) | .000 (-0.1) | .008 (-3.2) | -.04 (146.2) | 0.002 | .001 (-0.7) | .022 (-8.4) |
| | **Total gray** | Bil | -.05 (28.6) | 0.073 | .001 (0.8) | .002 (0.9) | .001 (-0.8) | .005 (-2.2) | -.05 (130.4) | 0.004 | .001 (-0.5) | .044 (-16.1) |
| | **Choroid plexus** | Left | -.05 (24.4) | -0.002 | .003 (1.2) | .001 (0.6) | .014 (5.0) | .011 (4.0) | -.03 (14.9) | -0.036 | .001 (0.8) | .005 (2.5) |
| | | Right | -.04 (16.7) | 0.002 | .000 (0.1) | .000 (0.3) | .022 (7.3) | .014 (4.8) | -.03 (14.5) | -0.037 | .001 (0.6) | .009 (4.0) |
| | **Optic chiasm** | Bil | -.01 (0.6) | -- | .000 (0.2) | .000 (0.1) | .005 (2.1) | .012 (4.4) | -.00 (0.1) | -- | .000 (-0.2) | .000 (-0.1) |
| | **Ventral DC** | Left | -.06 (30.2) | 0.014 | .000 (0.3) | .000 (0.2) | .000 (-0.3) | .003 (-1.3) | -.04 (9.0) | 0.000 | .002 (-1.3) | .001 (-0.9) |
| | | Right | -.06 (30.7) | 0.010 | .001 (-0.4) | .002 (-1.0) | .001 (-0.7) | .023 (-7.9) | -.05 (17.8) | -0.002 | .000 (-0.3) | .000 (-0.1) |
| | **Ventricle choroid** | Bil | -.06 (126.4) | -0.005 | .032 (12.1) | .016 (6.3) | .014 (5.6) | .008 (3.4) | -- | -- | -- | -- |
| | **Vessel** | Left | -.00 (0.2) | -- | .000 (0.2) | .000 (0.2) | .001 (-0.6) | .001 (-0.5) | .00 (0.1) | -- | .001 (0.4) | .002 (1.1) |
| | | Right | -.01 (1.9) | | .000 (-0.2) | .000 (-0.2) | .000 (0.0) | .000 (0.0) | .00 (0.0) | -- | .000 (-0.3) | .007 (-3.1) |
| **Intensity** | **3rd Ventricle** | Bil | -.05 (19.1) | 0.009 | .001 (0.8) | .001 (0.7) | .021 (-7.3) | .010 (-3.6) | -.03 (13.5) | -0.045 | .000 (0.2) | .001 (0.5) |
| | **4th Ventricle** | Bil | -.08 (23.6) | 0.026 | .000 (-0.3) | .000 (-0.3) | .007 (-2.7) | .001 (-0.5) | -.02 (2.0) | -- | .000 (0.0) | .003 (1.8) |
| | **5th Ventricle** | Bil | -.02 (0.5) | -- | .001 (0.7) | .002 (1.1) | .000 (-0.3) | .002 (-1.1) | .02 (0.5) | -- | .000 (0.1) | .005 (2.3) |
| | **Inf lat vent** | Left | -.05 (14.4) | -0.008 | .015 (-5.4) | .004 (-1.8) | .008 (-3.0) | .002 (-1.0) | -.05 (13.7) | -0.047 | .000 (-0.1) | .008 (-3.5) |
| | | Right | -.04 (8.4) | -0.003 | .002 (-1.0) | .001 (-0.4) | .002 (-0.8) | .000 (-0.1) | -.05 (11.3) | -0.044 | .000 (0.0) | .004 (1.8) |
| | **Lat vent** | Left | -.03 (4.5) | -0.007 | .000 (0.3) | .001 (0.5) | .007 (-2.6) | .000 (-0.0) | -.03 (6.5) | -0.081 | .000 (0.1) | .005 (2.4) |
| | | Right | -.03 (6.4) | -0.005 | .000 (0.1) | .000 (0.2) | .007 (-2.8) | .000 (-0.1) | -.03 (9.1) | -0.077 | .000 (0.2) | .006 (3.0) |
| | **CSF total** | Bil | -.05 (11.1) | 0.019 | .000 (0.2) | .000 (0.3) | .009 (-3.5) | .003 (-1.4) | -.04 (14.1) | -0.058 | .000 (0.0) | .010 (4.5) |
| | **Accumbens** | Left | -.10 (22.6) | -0.047 | .000 (0.1) | .000 (0.3) | .013 (4.5) | .038 (12.4) | .03 (3.0) | -- | .000 (-0.1) | .032 (-11.8) |
| | | Right | -.10 (22.8) | -0.055 | .000 (-0.1) | .000 (-0.0) | .007 (2.8) | .028 (9.5) | .04 (5.8) | -0.094 | .000 (0.0) | .043 (16.2) |
| | **Amygdala** | Left | -.03 (1.5) | -- | .000 (-0.1) | .000 (-0.0) | .001 (-0.5) | .002 (-0.8) | .04 (3.6) | -- | .000 (0.0) | .008 (3.4) |
| | | Right | -.03 (2.3) | -- | .001 (-0.4) | .001 (-0.6) | .001 (0.6) | .006 (2.4) | .04 (4.1) | -0.099 | .000 (0.0) | .008 (3.3) |
| | **Brainstem** | Bil | -.07 (7.0) | 0.058 | .000 (-0.1) | .000 (-0.3) | .000 (0.0) | .001 (0.8) | -.03 (1.3) | -- | .001 (-0.6) | .033 (-11.9) |
| | **Caudate** | Left | -.05 (8.0) | -0.025 | .000 (0.0) | .000 (0.2) | .002 (1.1) | .025 (8.4) | -.01 (0.7) | -- | .000 (0.0) | .003 (1.7) |

| | | | | | | | | | | | | |
|---|---|---|---|---|---|---|---|---|---|---|---|---|
| | | Right | -.05 (6.0) | -0.034 | .000 (-0.2) | .000 (-0.1) | .004 (1.7) | .024 (8.1) | .00 (0.2) | -- | .000 (-0.4) | .001 (-0.9) |
| | **Cerebellum** | Left | -.12 (34.8) | 0.041 | .000 (0.0) | .002 (0.8) | .000 (0.0) | .010 (3.6) | .03 (4.5) | -0.062 | .000 (0.2) | .003 (1.8) |
| | | Right | -.15 (47.9) | 0.025 | .001 (0.5) | .000 (0.1) | .001 (-0.7) | .000 (-0.2) | .00 (0.0) | -- | .000 (0.2) | .003 (1.7) |
| | **Hippocampus** | Left | -.03 (1.8) | -- | .000 (-0.2) | .000 (-0.2) | .006 (-2.4) | .001 (-0.6) | .02 (2.1) | -- | .001 (-0.6) | .014 (-5.5) |
| | | Right | -.07 (10.2) | 0.013 | .000 (-0.4) | .000 (-0.3) | .001 (-0.6) | .000 (-0.3) | .02 (1.0) | -- | .000 (0.0) | .004 (1.8) |
| | **Pallidum** | Left | -.00 (0.1) | -- | .001 (0.4) | .000 (0.0) | .001 (-0.6) | .009 (-3.3) | -.01 (0.3) | -- | .001 (0.8) | .011 (4.3) |
| | | Right | -.01 (0.4) | -- | .002 (1.2) | .000 (0.4) | .000 (0.0) | .002 (1.1) | .05 (5.2) | -0.007 | .001 (0.5) | .003 (1.7) |
| | **Putamen** | Left | -.03 (2.3) | -- | .001 (-0.5) | .000 (-0.1) | .004 (-1.6) | .001 (-0.6) | .01 (0.3) | -- | .000 (0.1) | .010 (4.2) |
| | | Right | -.00 (0.2) | -- | .000 (-0.1) | .000 (-0.0) | .000 (0.1) | .000 (0.3) | .04 (8.5) | -0.066 | .000 (0.3) | .000 (0.3) |
| | **Thalamus** | Left | -.06 (7.2) | 0.027 | .000 (-0.2) | .000 (-0.1) | .005 (-2.0) | .021 (-7.3) | -.01 (0.5) | -- | .000 (0.0) | .016 (5.9) |
| | | Right | -.06 (8.0) | 0.035 | .000 (-0.1) | .000 (-0.1) | .007 (-2.8) | .017 (-5.8) | .01 (0.3) | -- | .001 (-0.5) | .010 (-4.0) |
| | **CC anterior** | Bil | -.04 (5.5) | 0.017 | .006 (-2.4) | .006 (-2.4) | .001 (0.4) | .004 (1.7) | -.03 (3.5) | -- | .000 (0.3) | .015 (5.8) |
| | **CC central** | Bil | -.04 (4.6) | 0.024 | .000 (-0.1) | .000 (-0.3) | .000 (-0.3) | .000 (-0.0) | -.03 (2.1) | -- | .001 (0.5) | .025 (9.0) |
| | **CC mid anterior** | Bil | -.03 (3.3) | -- | .001 (-0.5) | .000 (-0.2) | .000 (0.0) | .001 (0.6) | -.04 (4.0) | 0.090 | .000 (-0.3) | .020 (-7.6) |
| | **CC mid posterior** | Bil | -.06 (22.1) | 0.025 | .001 (0.7) | .000 (0.3) | .006 (-2.5) | .009 (-3.3) | -.03 (3.8) | -- | .002 (1.4) | .031 (11.5) |
| | **CC posterior** | Bil | -.04 (8.2) | 0.031 | .000 (0.1) | .000 (0.2) | .000 (0.3) | .000 (0.0) | -.02 (0.9) | -- | .001 (-0.5) | .000 (-0.1) |
| | **Cerebellum WM** | Left | -.11 (12.8) | 0.097 | .000 (-0.1) | .000 (-0.2) | .008 (-3.2) | .009 (-3.3) | -.01 (0.3) | -- | .000 (0.1) | .000 (0.3) |
| | | Right | -.10 (12.2) | 0.062 | .000 (-0.2) | .000 (-0.1) | .013 (-4.7) | .003 (-1.4) | -.01 (0.3) | -- | .001 (-0.5) | .000 (-0.4) |
| | **WM hypointensities** | Bil | -.02 (1.7) | -- | .000 (0.1) | .001 (0.5) | .001 (-0.7) | .022 (-7.5) | -.06 (23.2) | 0.000 | .001 (-0.3) | .001 (-0.4) |
| | **Non WM hypointensities** | Bil | -.00 (0.0) | -- | .000 (-0.1) | .000 (-0.0) | .000 (0.4) | .003 (1.2) | .01 (0.3) | -- | .000 (0.1) | .005 (2.3) |
| | **Choroid plexus** | Left | -.06 (25.1) | 0.011 | .000 (-0.1) | .000 (-0.0) | .018 (-6.4) | .012 (-4.4) | -.05 (32.9) | -0.003 | .001 (-0.5) | .001 (-0.6) |
| | | Right | -.05 (15.3) | -0.009 | .000 (0.0) | .001 (0.6) | .032 (-10.7) | .014 (-5.0) | -.06 (37.9) | -0.009 | .001 (-0.8) | .005 (-2.2) |
| | **Optic chiasm** | Bil | -.01 (0.5) | -- | .000 (-0.1) | .003 (-1.4) | .000 (-0.1) | .000 (-0.4) | -.01 (0.3) | -- | .000 (-0.4) | .004 (-1.9) |
| | **Ventral DC** | Left | -.02 (0.9) | -- | .001 (0.6) | .000 (0.2) | .000 (-0.4) | .001 (-0.4) | -.01 (0.2) | -- | .000 (-0.1) | .028 (-10.1) |

| | | | | | | | | | | | | |
|---|---|---|---|---|---|---|---|---|---|---|---|---|
| | | Right | -.02 (0.6) | -- | .001 (0.4) | .000 (0.0) | .000 (0.0) | .003 (1.3) | .00 (0.0) | -- | .000 (-0.1) | .018 (-6.8) |
| | **Vessel** | Left | -.01 (0.4) | -- | .000 (0.1) | .001 (0.5) | .004 (-1.9) | .000 (-0.1) | -.01 (0.5) | -- | .000 (-0.3) | .006 (-2.6) |
| | | Right | -.02 (0.7) | -- | .000 (0.0) | .000 (0.1) | .002 (-0.8) | .001 (-0.5) | -.02 (2.8) | -- | .000 (0.1) | .017 (6.4) |

**Supplementary Table 4. Relationship between *brain age delta* and change in brain features.** Long. change = Longitudinal change in the *raw* neuroimaging features (mean change [$\log_{10}$(p)]). PC1 load = feature loadings on the first component of longitudinal change. $Delta_{cross}$ = Relationship between cross-sectional *brain age delta* and feature change ($r^2$ [$\log_{10}$(p)]). $Delta_{long}$ = Relationship between longitudinal *brain age delta* and feature change ($r^2$ [$\log_{10}$(p)]). GWC = Gray-white matter contrast. Cth = Cortical Thickness. Bil = Bilateral. Subc = Subcortical. n = 1372 and 1500 for the UK Biobank and the Lifebrain datasets. |N| = 365 AND 372 features in the UK Biobank and the Lifebrain datasets. XGB = boosting gradient as implemented in XGBoost.

| Lifebrain Consortium (http://www.lifebrain.uio.no/about/) | | | | |
|---|---|---|---|---|
| **LCBC** | http://www.oslobrains.no | Kristine B. Walhovd | | Norwegian Regional Committee for Medical and Health Research Ethic; Regional Ethical Committee of South Norway |
| **BETULA** | http://www.ufbi.umu.se/english | Lars Nyberg | | Regional Ethical Vetting Board at Umeå University |
| **BASE-II** | https://www.mpib-berlin.mpg.de/en/research/lifespan-psychology | Ulman Lindenberger | | Ethics committee of the Charité-Universitätsmedizin *Berlin* |
| **Cam-CAN** | https://www.cam-can.org/ | Lorraine K. Tyler & Richard Henson | & | Cambridgeshire 2 Research Ethics Committee |
| **UB** | http://www.ub.edu/bbslab/bbslab/ | David Bartrés-Faz | | Comisión de Bioética de la Universidad de Barcelona and Hospital Clinic |
| **AIBL*** | https://aibl.csiro.au/research/ | Christopher Rowe | | Institutional ethics committees of Austin Health, StVincent's Health, Hollywood Private Hospital and Edith Cowan University |

**Supplementary Table 5. Contact information.** Contact information and ethical comittees for the different cohorts of the Lifebrain consortium.

*AIBL does not belong to the Lifebrain consortium.

| Sample | Scanner | Sequence | Tesla | Slices | Voxel size (mm) | Time parameters (TR / TE / TI [ms]) | Other parameters (FA / FOV [∘/mm]) |
|---|---|---|---|---|---|---|---|
| **UK Biobank (main sample)** | | | | | | | |
| UK Biobank* | Skyra Siemens | 3D MP-RAGE | 3.0 | 256 | 1 x 1 x 1 | 2,000/-/880 | |
| **Lifebrain (replication sample)** | | | | | | | |
| LCBC | Avanto Siemens | 3D MP-RAGE | 1.5 | 160 | 1.25 x 1.25 x 1.25 | 2,400/3.61/1,000 | 8/240x240 |
| | Skyra Siemens | 3D MP-RAGE | 3.0 | 176 | 1 x 1 x 1 | 2,300/2.98/850 | 8/256 x 256 |
| | Prisma Siemens | 3D MP-RAGE | 3.0 | 208 | 1 x 1 x 1 | 2,400/2.22/1,000 | 8/240 x 256 |
| Cam-CAN | Tim Trio Siemens | 3D MP-RAGE | 3.0 | 192 | 1 x 1 x 1 | 2,250/2.98/900 | 9/256 x 240 |
| Base-II | Tim Trio Siemens | 3D MP-RAGE | 3.0 | 176 | 1 x 1 x 1 | 2,500/4.77/1,100 | 7/256 x 256 |
| Betula | Discovery GE | 3D FSPGR | 3.0 | 176 | 1 x 1 x 1 | 8.19/3.2/450 | 12/250 x 250 |
| UB | Tim Trio Siemens | 3D MP-RAGE | 3.0 | 240 | 1 x 1 x 1 | 2,300/2.98/900 | 9/256 x 256 |
| AIBL** | Avanto Siemens | 3D MP-RAGE | 1.5 | 160 | 1 x 1 x 1.2 | 2,300/2.98/900 | 9/240 x 256 |
| | Verio Siemens | 3D MP-RAGE | 3.0 | 160 | 1 x 1 x 1.2 | 2,300/2.98/900 | 9/240 x 256 |
| | Tim Trio Siemens | 3D MP-RAGE | 3.0 | 160 | 1 x 1 x 1.2 | 2,300/2.98/900 | 9/240 x 256 |

**Supplementary Table 6. Data acquisition parameters.** Data acquisition parameters for the T1w sequences**.** *UK Biobank employed three scanners of the same model and with equivalent parameters (Cheadle, Reading, and Newcastle centers). **AIBL does not belong to the Lifebrain consortium but was included in the Lifebrain replication dataset.
